# Ultimate paths of least resistance: Intrinsically disordered links as developmental resets in regulatory protein networks

**DOI:** 10.1101/2025.03.17.643830

**Authors:** Alexander V. Badyaev, Carmen Sánchez Moreno, Cody A. Lee, Sarah E. Britton, Laurel M. Johnstone, Renée A. Duckworth

## Abstract

Development and evolution require both stability and adaptability, yet how these opposite properties are reconciled is unclear. Here, we show that intrinsically disordered proteins (IDPs) act as reset mechanisms in conserved regulatory networks facilitating developmental transitions by integrating physical processes with genetic regulation. By tracing the ontogeny of mesenchymal cells in avian beak primordia, we demonstrate that mechanosensitive IDPs mediate shifts between physical cell states via dosage-dependent binding plasticity, converting stochastic protein variation into discreet regulatory networks. The disorder-enabled connectivity in these proteins resets their regulatory specialization and promotes population divergence. Comparative analyses across avian proteomes confirm that binding plasticity in transcriptional IDPs drives their diverse regulatory associations and accelerates their evolution. By resetting specialized states in conserved regulatory networks, IDPs flexibly regulate developmental pathways and reconcile precision with evolvability.

---

Sequential differentiation in development and adaptations in evolution must necessarily negotiate a tension between specialization and continuity, such that the strength and precision of specialized states does not interfere with subsequent change. Because of this tension, regulators of specialized states are rarely state-specific themselves. Instead, evolution is thought to favor weak and reversible regulators that are able to crystallize or absorb existing variation to either prime transitions or maintain uniformity within specialized states^1^. To have such effects, these regulators must be easily recognized by their targets and be able to rapidly assemble and disassemble regulatory complexes to match specific responses to external contexts^2-5^.

In development, these contrasting demands of generality and context-specificity are met by evolutionarily conserved transcriptional organizers that accumulate a large repertoire of conditionally responsive factors^6,7,^ (exemplified by Mediator complex)^8,9^. Although this organizational principle is central in molecular regulation, the mechanisms by which these complexes reconcile generality of their content with precision of its deployment are not fully understood^10^. An emergent hypothesis is that this ability is conferred on regulatory proteins by the extent and juxtaposition of ordered and intrinsically disordered regions (IDR) in their structural conformations that give these intrinsically disordered proteins (IDP) the ability to integrate general physical responsiveness and biological specificity in orchestrating regulatory networks^11-14^.

Four mechanisms are central for IDPs’ proposed role in reconciling generality and context-specificity in molecular regulation. First is their propensity to induce physical phase-separation and form droplet-like membraneless organelles or nuclear condensates into which they accumulate a multitude of conditionally responsive regulators^15-19^. These complexes can then persist across developmental stages and even generations^20^, until the external context and their stored conditionalities match. The sequestration of these regulators into condensates is facilitated by the second key property of IDPs – their dosage-dependent binding plasticity, a propensity to bind to the most locally abundant or most proximate partners regardless of binding specificity^21-23^ a property that is facilitated by induced folding^24-27^. Further, conformational ensembles of IDPs behave as material states – they have elasticity, volume and reach – accounting for their ability to bridge physical and signaling processes governing cell behavior^17,22,23,28-34^. Finally, juxtaposition of ordered and disordered regions within a protein enables their mutual regulation; IDPs can activate or suppress their own activity without binding to external regulators^35^. This combination of regulatory autonomy and bidirectional fluency with general physical and specific biological contexts should make IDPs universal organizers of regulatory networks able to match specific biological signaling to a range of developmental contexts^36-38^.

Here we examine whether IDP-enabled organization, where conserved transcriptional regulators capitalize on their intrinsic disorder to accumulate an array of conditionality-sensitive factors (Fig. 1), accounts for evolutionary persistence and functional versatility of a core regulatory network underlying avian beak development^39-41^. This network is closely tuned to cellular and histological processes throughout development, forming topologically plastic configurations with context-dependent effects on cellular dynamics (Fig.1a, Supplementary Text). Yet, it persists largely unchanged over evolutionary time, underlying both deep evolutionary conservation and ecological-scale plasticity in beak morphology^42-47^. IDP-enabled organization could reconcile this conservation with context-specificity of its outcomes, explaining also why these ubiquitous transcriptional regulators rarely define regulatory network-to-phenotype maps^48-50^, but instead make these maps possible, preserving the network cohesion across specialized states^51,52^. We envision three broad scenarios for the role of IDPs in persistence of this network (Fig. 1d).

**Fig. 1.**
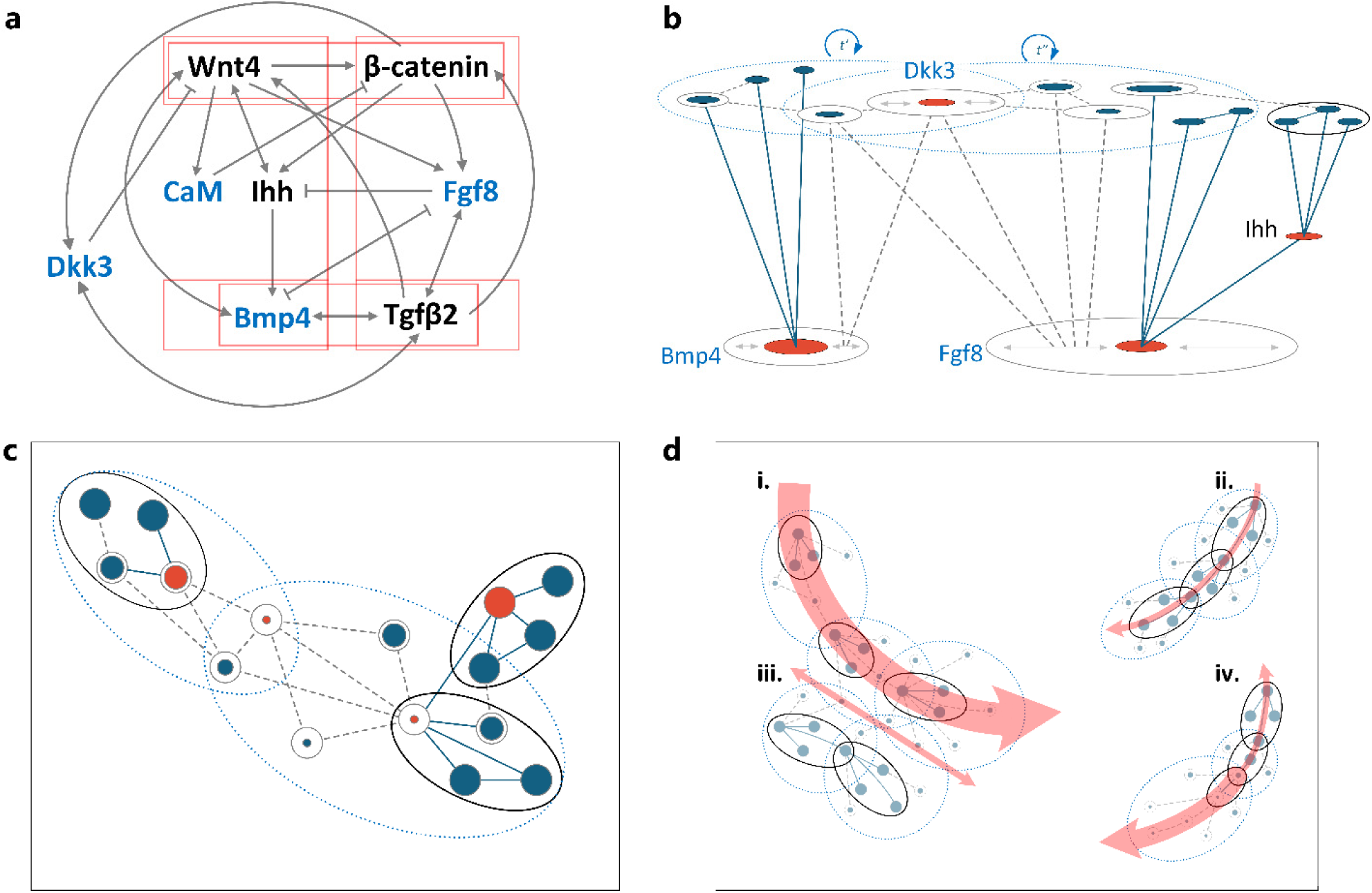
Evolutionary consequences of integration between a network of intrinsically disordered transcriptional regulators and morphogenetic processes. **(a)** The network of key proteins central for beak development in this study (based on Supplementary Text). Arrows show upregulation; bars – downregulation. Red boxes enclose conserved signaling motifs; proteins in blue are intrinsically disordered (ID) (Supplementary Table S7). **(b)** Expansion of the key regulatory network orchestrated by IDP’s dosage-dependent binding and phase-separation of resulting protein complexes. Double headed arrows within each ellipsoid (filled red area shows protein concentration) reflect the ID range and associated dosage-dependent binding plasticity. Dashed lines connecting proteins indicate non-specific disorder-enabled binding; solid lines show specific, dosage-independent bindings. Four transcriptional regulators [from (a)] are shown for illustration. Note that some elements of the key network in (a) have only disorder-enabled links (e.g., “Dkk3”), coexpression of proteins can be caused by both direct binding and through binding intermediaries, and disorder-enabled partnering can produce both up- and down-regulating effects. Uncharacterized proteins (dark blue circles) are recruited by the known core IDPs into liquid-state condensates (blue dashed ellipsoids). Circular arrows with *t* indicate that these condensates can persist across developmental contexts. **(c)** Regulatory landscape of (b) shown in one plane (“view from above”). Solid ellipsoids indicate network modules unaffected by disorder-enabled connectivity; blue dashed ellipsoids illustrate the network expansion through the disorder-enabled connectivity. All other symbols as in (b). **(d)** Evolutionary scenarios of IDP-enabled integration of regulatory network and morphogenesis. Red transparent arrows show direction and width of an evolutionary path as shaped by regulatory network connectivity. Scenario ***i***: evolution proceeds through bridges formed by IDPs’ connectivity, ***ii***: IDP-enabled connectivity buffers evolutionary change that proceeds only through functional modules, ***iii***: evolutionary change is channeled by functional modules and enabled by IDPs’ stochastic connectivity, ***iv:*** disorder and associated binding plasticity is a temporary evolutionary stage.

First, disorder-enabled IDP connectivity may form temporary bridges between conserved regulatory network modules, facilitating their recombination (scenario *i,* Fig. 1d) or buffer and shield functional modules, maintaining plasticity around their evolutionary trajectory (scenario *ii,* Fig. 1d). Second, some disorder-enabled connectivity may be stochastic^53^, introducing developmental noise that channels evolutionary change orthogonally to functional modules (scenario *iii*, Fig. 1d). Finally, disorder and associated binding plasticity may be a transient evolutionary stage, diminishing when favorable connectivity patterns stabilize, increasing during functional complex breakdown, or undergoing disorder-to-order transitions upon binding (scenario *iv*, Fig. 1d). These scenarios assume distinct selection pressures on disordered linkers and ordered domains within IDPs, predicting contrasting patterns of IDP sequence divergence^23,54-56^.

Here we examine these scenarios for the role of IDPs in maintenance of a regulatory network at three levels: early-developmental transitions between mechanical cell states, among recently diverged populations, and across species. We first investigated aspects of regulatory network structure tuned to cell jamming—a ubiquitous transition between liquid-like and solid-like physical phases where homogeneous cell groups spontaneously form and dissolve small islands of densely packed (“caged”) slow-moving cells^57-60^. Although this transition is an essential feature of early multicellular development, it is invariant to local adaptations and exemplifies the most basic integration of physical processes driving jamming with biological processes by which cells modulate them^61-63^. Consequently, network features enabling cell jamming transitions reflect the continuity of early embryonic development prior to the onset of tissue differentiation and modification. To assess whether compensatory mechanisms enabling local transitions during early development are retained over evolutionary time, we first examine evolutionary divergence in this regulatory network across recently diverged house finches (*Haemorhous mexicanus*) populations^64^ and then explore prevalence of these mechanism across bird species (Fig. 1d).

We developed high throughput analytical tools enabling us to follow ontogenetic rearrangements and signaling dynamics in hundreds of thousands of mesenchymal cells across the upper beak primordia^65^ and found that the connectivity of intrinsically disordered transcriptional regulators is tuned to cell jamming transitions. The axes of the core regulatory network governing shifts between mechanical cell states align with patterns of evolutionary divergence in regulatory networks, a pattern driven by dosage-dependent binding plasticity of IDPs. Thus, intrinsically disordered links in regulatory networks function as developmental resets, enabling reversible transitions between specialized states. By integrating mechanosensitivity with regulatory plasticity, these proteins facilitate both developmental precision and evolvability, as exemplified in avian beak morphogenesis.

## Results

### Intrinsic disorder conveys mechanosensitivity to transcriptional regulators

Summarizing morphological variation of *n* = 724,465 mesenchymal cells into multivariate measures of cell shape (Supplementary Tables S2a), cell uniformity within groups (Supplementary Tables S3b), and jamming state (Supplementary Figs. S1-S2) we found close association between these measures and protein expression (Fig. 2). For example, when cells in a group became jammed, the spread (the proportion of cells expressing the protein, Methods) of Dkk3 and Bmp4 markedly contracts, whereas the spread of β-catenin, Calm1 and Fgf8 expands (Supplementary Table S4). Expression of Ihh and Tgfβ2 was not affected by cell jamming, but covaried with cell shape and uniformity (Fig. 2b,c).

**Fig. 2.**
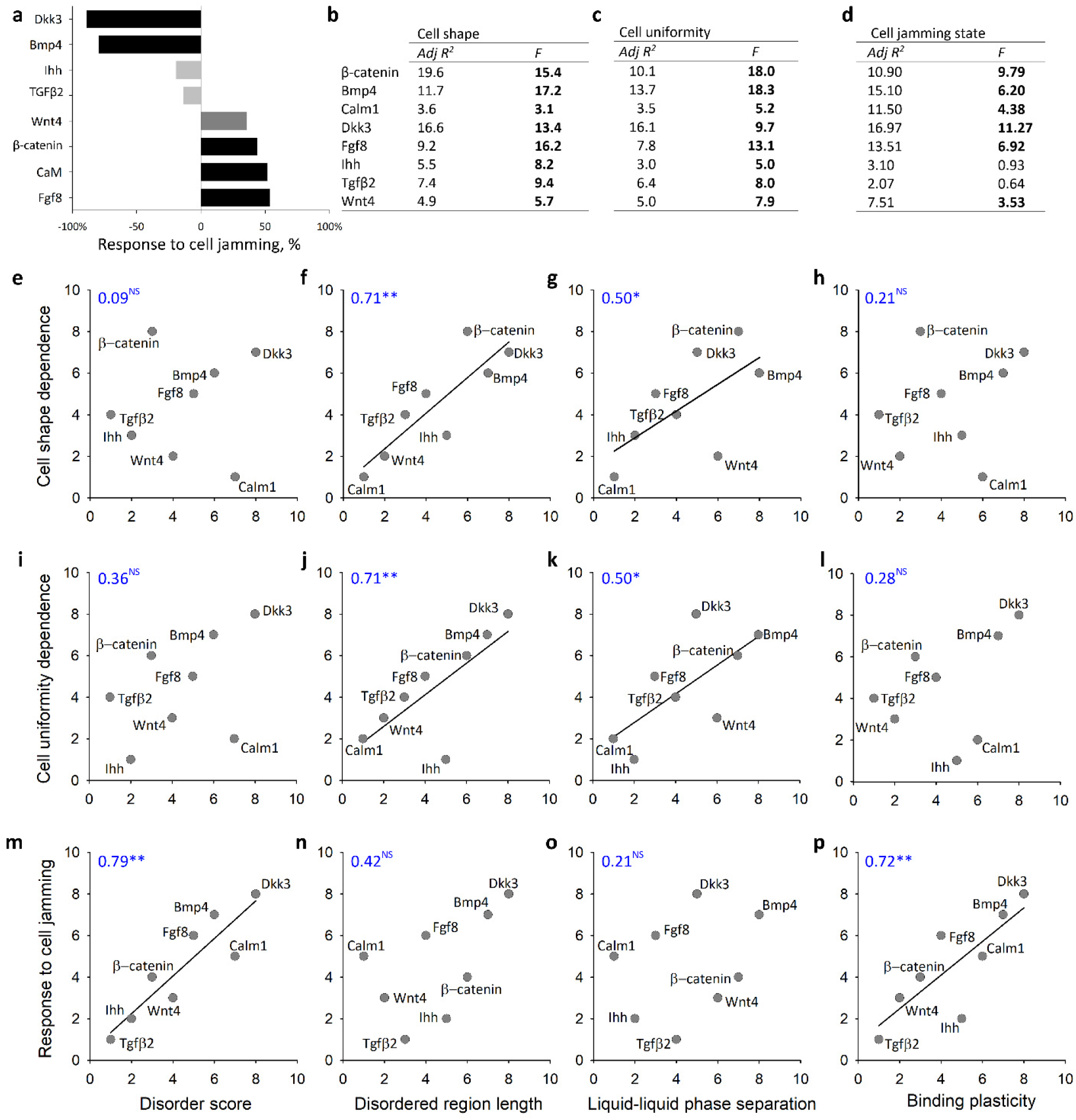
Mechanosensitivity of regulatory proteins covaries with their intrinsic disorder. **(a)** Change in protein expression between jammed and unjammed cell groups. Bars are relative change (%) between the least squared means for jammed and unjammed cell states. Black bars: *P* < 0.01, dark gray bars: *P* < 0.05, non-significant – light gray bar (data in Supplementary Table S4). Mixed-effects model of dependency of protein expression on **(b)** cell shape (Supplementary Table S2a-c), **(c)** cell uniformity (Supplementary Table S3a-c), and **(d)** cell jamming state (Supplementary Table S4). Each table shows residuals controlling the effects of population, developmental stage and their interaction (Supplementary Table S2b, S3b). Bold *F*-values are significant at *P* < 0.05 after Sidak adjustment for multiple comparisons. **(e-p)** Protein’s intrinsic disorder and phase separation propensity underlie correlation of its expression with **(f)** cell shape (Supplementary Table S2ab) and **(j)** cell uniformity within a group (Supplementary Table S3ab). Expression of proteins with **(m)** greater disorder and **(p)** greater binding plasticity are more sensitive to cell jamming transitions (Supplementary Table S4). Shown are Kendall’s correlational coefficients on ranks of Adj R2 contribution (b-d) and protein disorder properties (Supplementary Table S7).

To examine whether mechanosensitivity of these proteins (Fig. 2) is associated with their intrinsic disorder, we obtained their amino acid sequences (Supplementary Data S2) from whole genome sequencing of 20 house finches across the studied populations (Supplementary Table S1, Methods), and evaluated the proteins’ structural conformation, location and extent of intrinsic disorder and its biophysical consequences, as well as binding plasticity (Methods, Supplementary Figs. S3-S11, Supplementary Table S7). We find that more disordered proteins are more responsive to changes in cell shape, tissue uniformity, and cell jamming (Fig. 2). Proteins with greater IDRs and a greater propensity for phase separation are most sensitive to variation in cell shape and cell uniformity within a group (Fig. 2 f, j, g, k), whereas more disordered proteins with greater binding plasticity respond the strongest to cell jamming transitions (Fig. 2 m, p). This suggests that the IDPs’ mechanosensitivity is mediated by their ability to form or maintain binding with other proteins under changing physical conditions. We thus examined the link between protein mechanochemistry and binding plasticity directly.

### Dosage-dependent binding plasticity orchestrates distinct regulatory networks under cell jamming

Dosage-dependence of binding plasticity predicts that networks involving IDPs will have distinct topologies across contexts with variable protein concentrations Fig. 1b, ^21^. Indeed, we find repeatable differences in protein networks between jammed and unjammed cell states (Fig. 3a,b, Supplementary Table S5ab) which differ in protein expression (Fig. 2). With the exception of Wnt4 (which, although having a long IDR in the house finch (Supplementary Table S7), does not reach the 30% threshold for conventional classification as an IDP, Methods), the differences between the two coexpression networks are due to the gains and losses of IDP partners: changes in connectivity of Calm1, Wnt4, Dkk3, and Bmp4 contribute the most to the regulatory network divergence (Fig. 3f, Supplementary Table S5c). For example, in the jammed state, Dkk3 retains just two (Calm1 and Ihh) of the seven coexpression partners it has in the unjammed state. Similarly, in the jammed state, disordered Bmp4 loses all six coexpression partners it has in the unjammed state but gains a new coexpression link to Calm1, which, in turn, gains two more coexpression partners in the jammed state compared to the unjammed state (Fig. 3a,b). In these examples, disordered Dkk3 and Calm1 connect network motifs (Fig. 1a, Supplementary Text), such that their mechanosensitivity would allow the rearrangement of these modules in different contexts (Fig. 3f). To examine the evolutionary persistence of these patterns, we next examine the stability of regulatory networks across populations and the maintenance of structural disorder in IDPs in relation to their amino acid sequence divergence across species.

**Fig. 3.**
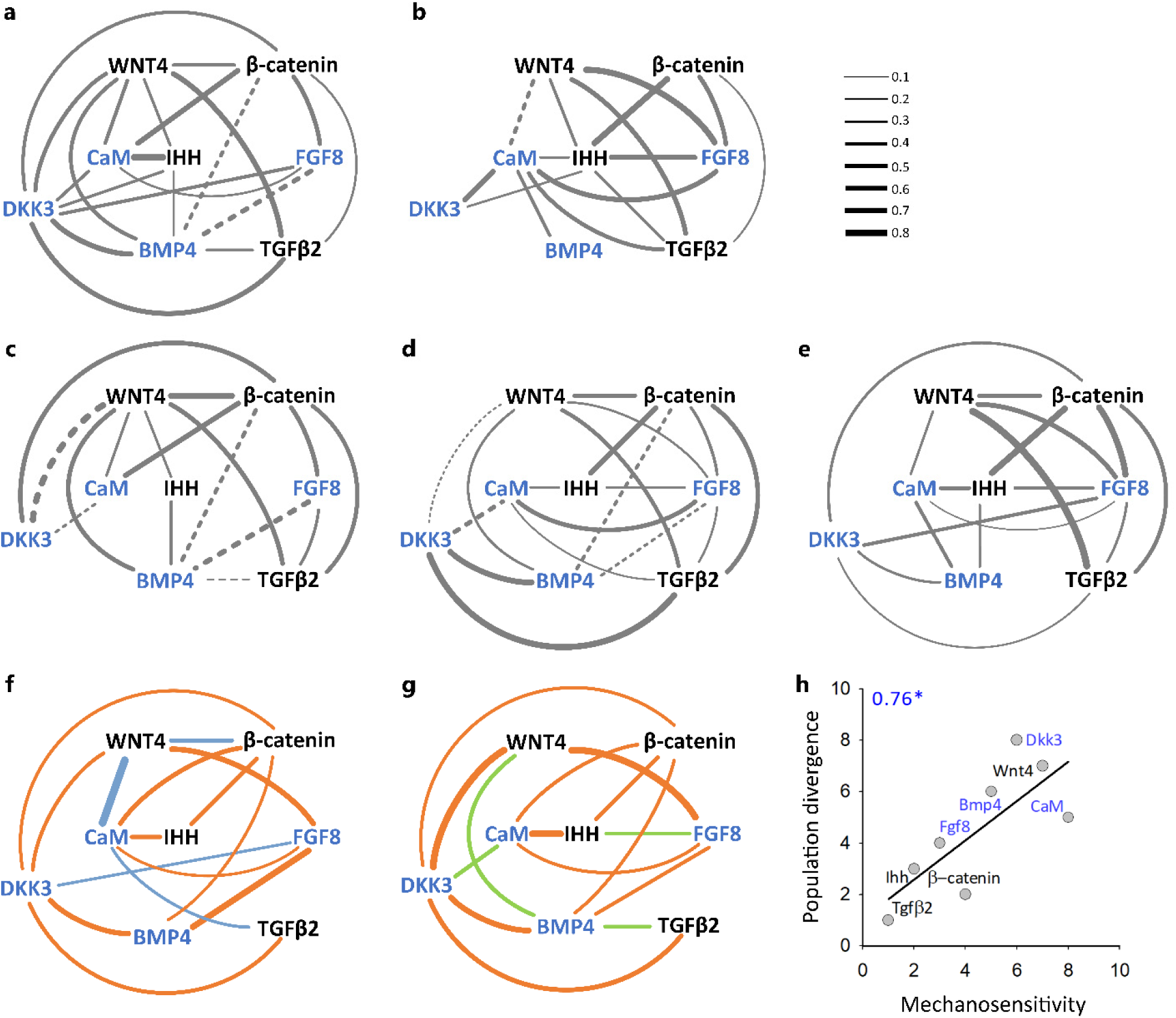
The axes of mechanosensitivity are the axes of population divergence in regulatory networks and both are delineated by disorder-induced connectivity of IDPs (proteins shown in blue). **(a)** Unjammed and **(b)** jammed cell states. Line thickness shows strength of partial correlations, dashed lines show anticorrelation. Only significant partial correlation coefficients are shown (based on Supplementary Table S5 a,b). Correlated expression of proteins in **(c)** nMT populations (Supplementary Table S6a), **(d)** eMT populations (Supplementary Table S6b), and **(e)** wMT populations (Supplementary Table S6c). **(f)** Visualization of divergence matrix of jammed *vs*. unjammed cell states (Supplementary Table S5c) and **(g)** visualization of population divergence matrix (Supplementary Table S6d). Only greater than average divergence links are shown (see Supplementary Tables S6 for details and derivation). Links in orange are shared between cell state and population divergence matrices. Links in blue and green are unique to cell state and population divergence matrices, respectively. **(h)** Contribution of individual proteins to divergence matrix of the cell state sensitivity (**f**, Supplementary Table S5c) in relation to contribution of this protein to the population divergence matrix (**g**, Supplementary Table S6g) and associated Kendall’s correlation coefficient.

### The axes of path-resetting between cell mechanistic states are the axes of evolutionary divergence

We first derive protein coexpression networks for northern, eastern, and western Montana populations (Fig. 3c-e, Supplementary Table S1, S6a-c) and find that the populations diverged mostly in IDPs’ coexpression (Fig. 3g): Dkk3, Wnt4, Bmp4, and Calm1 contribute the most to population divergence (Supplementary Table S5d-g). For example, coexpression of Dkk3 and Tgfβ2 is strong in eastern Montana, weak in central Montana and not detectable in northern Montana. Similarly, the antagonistic coexpression between Fgf8 and Bmp4 is present in northern and eastern, but not in western Montana (Fig. 3c-e, Supplementary Table S5d-f). Patterns of connectivity in the cell state divergence matrix (Fig. 3f) and population divergence matrix (Fig. 3g) are remarkably congruent (Fig. 3h) and formed by the same IDPs (Supplementary Table S6h).

### Binding plasticity of IDPs underlies state transitions

Having established that IDP connectivity delineates the population divergence of regulatory networks, we examined likely evolutionary scenarios behind this pattern (Fig. 1d). Scenario *i* (Fig. 1d) would lead to weaker or fluctuating selection on binding specialization, driving the greater sequence divergence and faster evolution of IDPs compared to ordered proteins. Under scenario *ii* (Fig. 1d), IDPs or IDRs can be under stabilizing selection to preserve network cohesion leading to conservation of disorder properties or to opposite selection pressures between disordered linkers and stable domains and their coevolution. Scenario *iii* also predicts undiminished disorder properties over evolutionary time and channeling of developmental variability by regulatory modules. Finally, scenario *iv* predicts rapid evolution of disorder properties, including periodic shifts between order and disorder (Fig. 1d).

To test these predictions, we first calculated the disorder properties of focal proteins across all available avian proteomes (Methods, Fig. 4, *n* = 70-170 species, Supplementary Tables S7-8, Supplementary Data S3-11). To directly compare the evolutionary divergence between these proteins, we further extracted data for *n* = 57 species that had complete data of comparable quality for all the focal proteins (Methods, Supplementary Table S9). We find that for each protein, the extent of disorder is largely conserved across species (Fig. 4). However, the propensity to form phase separation varies extensively across species in Dkk3, Bmp4, and Wnt4, without corresponding changes in the overall disorder score (Fig. 4a, e, i), corroborating findings that the propensity to phase-separate is not associated with the overall disorder across species (Fig. 5b). More disordered proteins – e.g., highly disordered Calm1, Calm2, and Bmp4 – have greater binding plasticity (Fig. 5a). In turn, proteins with greater binding plasticity and, correspondingly a wider range of binding partners, accumulate greater amino acid sequence divergence and have more variable evolutionary rates than proteins with higher binding specialization (Fig. 5c, d).

**Fig. 4.**
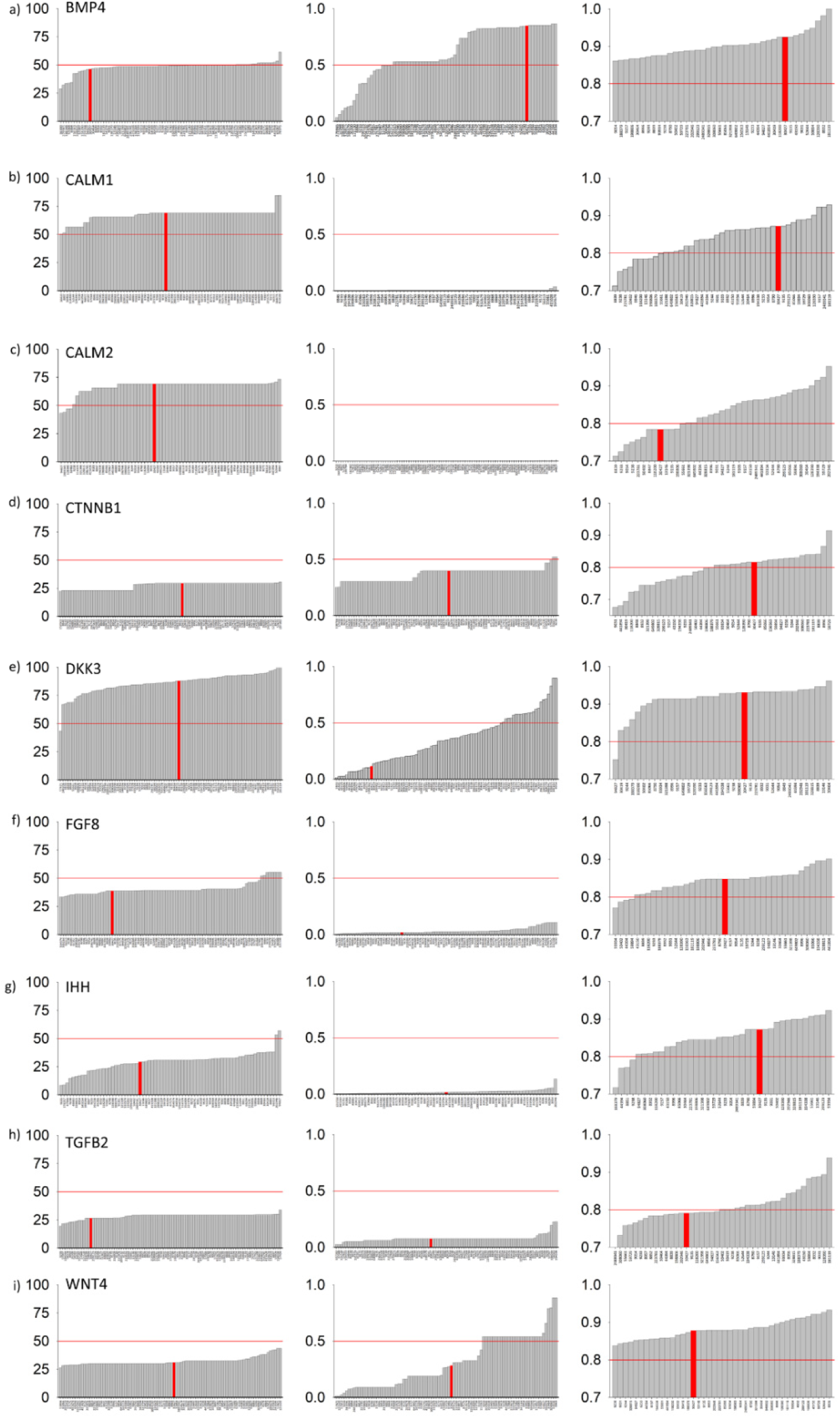
The variable physical arrangements of amino acids in IDRs of conserved protein sequences affect their biophysical properties. Disorder score, % (left column), propensity to form liquid-liquid phase separation (center) and binding affinity (right) for **a)** Bmp4 (*n* = 120 species), **b)** Calm1 (*n* = 70 species), **c)** Calm2 (*n* = 77 species), **d)** CTNNB1 (β catenin; *n* = 170 species), **e)** Dkk3 (*n* = 103 species), **f)** Fgf8 (*n* = 122 species), **g)** Ihh (*n* = 72 species), **h)** Tgfβ2 (*n* = 100 species), and **i)** Wnt4 (*n* = 119 species) arranged in the order of increase (based on Supplementary Table S7). Values above the red lines indicate species whose protein versions are mostly disordered, able to form phase separation and have higher than average binding affinity. Red bars are values for the house finch proteins in this study (Supplementary Data 2).

**Fig. 5.**
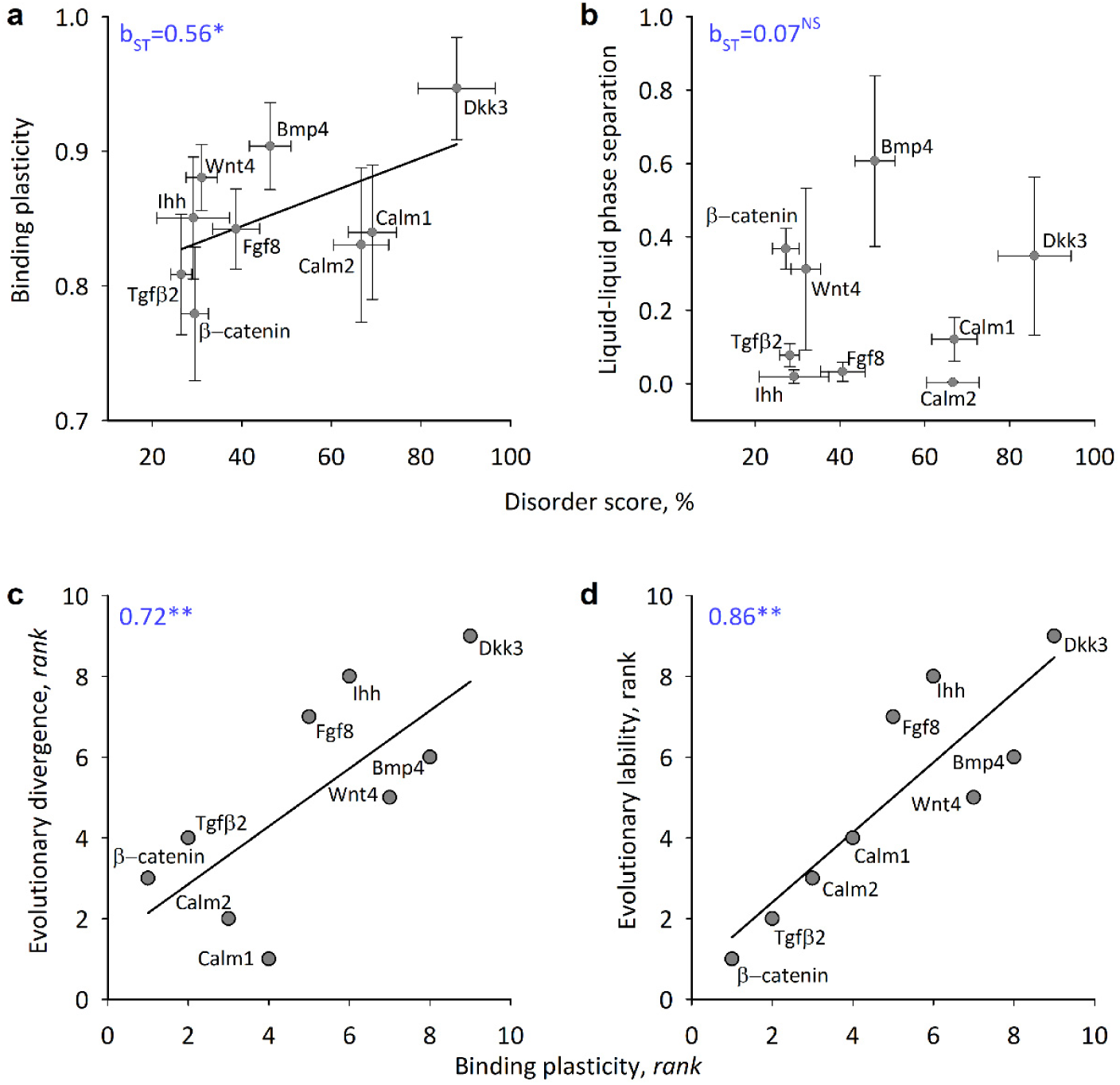
Binding plasticity of IDPs underlies their evolutionary divergence. **(a)** Disorder score correlates with binding plasticity, but not with **(b)** phase separation propensity in proteins across bird species (Supplementary Table S7). Shown are means and s.d., and standardized regression coefficients. Greater binding plasticity and corresponding diversity of binding partners and contexts (Fig. 4) is associated with **(c)** faster and **(d)** more variable evolutionary divergence. (c) mean and (d) s.d. of divergence among percent identity matrices of aligned sequences across *n* = 57 species (Methods, Supplementary Tables S7-S9). Kendall correlation coefficients are shown.

## Discussion

In the context of the dynamic and continuous process of evolution, specialization must necessarily be reversible. Thus, the historical tension between the strength and precision of adaptation and the ability to change it^66-68^ has shifted to a debate on whether this ability evolves specifically, controlled by dedicated master switches, or is an unescapable byproduct of a dynamic and multilevel developmental process^1,5,49,69-73^.

We found that the same axes of a core regulatory network that enable ubiquitous transitions between cell mechanical states also delineate its evolutionary divergence (Fig. 3h). These axes are formed by dosage-dependent binding that converts stochastic variability in protein expression into repeatable regulatory networks, essentially delineating the space of subsequent exploration and divergence at the population level (Fig. 3c-e, Supplementary Table 6h). Notably, this pattern occurs in fields of homogeneous cells at developmental stages that precede population-specific tissue modification (e.g., prior to beak condensation formation). Thus, rather than regulating specialized states, these axes essentially reset path-dependency (historicity) between consecutive states, assuring developmental continuity.

These results offer an alternative explanation for the often-documented alignment of variation produced by distinct intrinsic developmental sources (e.g., mutation- or stress-induced developmental variation, standing genetic variance, phenotypic noise)^74-78^. This alignment is thought to be produced by the developmental architecture that has coevolved with patterns of functional and genetic integration or by variational properties of genetic regulation^79-85^. If the dosage-dependent connectivity that delineates path-resetting directions in this study (Fig. 3g) is produced by IDPs’ conformational ensembles (e.g., by induced folding), then these links are shielded from mutational input (see below). This decoupling of mutational input from structural conformation and of binding strength from binding specificity makes these links the ultimate reset mechanisms, underlying their propensity to channel developmental variability (Fig. 1a, scenario *iii*). This maintains the directions of variability even in increasingly complex phenotypes, assures continuity of development and may account for the alignment of genetic and phenotypic developmental variance on a macroevolutionary time scale^52,86,87^.

In biology, unlike human engineering, the specialization of an organism only emerges in the context of interaction with other coevolving partners; it is neither preexisting nor persisting beyond the interaction. Thus, the central tension in debates on biological specialization versus evolutionary change is not the strength of specialization per se, but the ability to reset past dependencies between the coevolutionary partners^1,88^. On the most basic level, the integration of any processes not sharing an evolutionary history leads to the resetting of the evolutionary trajectory of specialization. In this sense, fluent bidirectional transitions in regulatory IDPs between path-independent (i.e., ahistoric) physical processes and highly contingent biological processes (whose impact requires coevolution) illustrate how specialization and changeability is combined in biology. For example, many IDPs undergo a disorder-to-order transition upon binding but maintain a disordered state when not bound; the coupling of folding and binding allows some binding partners to maintain a disordered state even when bound^4,26,89^. These properties lessen mutually imposed selection between binding partners and reconcile binding specialization with undiminished binding plasticity and thus an opportunity for change^29,30^. The coevolution and the associated compensatory relationship between ordered and disordered regions within IDPs (conformational buffering) offers another mechanistic illustration. In conformational buffering, a protein’s functionality is maintained despite environmental fluctuations, mutations, or changes in binding partners, because disordered linkers can compensate for mutations in ordered motifs, whereas ordered motifs can offset the physical forces acting on disordered linkers^23^. The coevolution of disordered and ordered regions can transfer the effect of selection and environmental pressures to partners that are not typically direct targets – e.g., disordered regions can be under functional selection when they coevolve with ordered motifs, and ordered regions can become sensitive to the physicochemical context of cells or tissues when it affects their disordered partners^22,30,90^.

IDPs are thought to evolve faster than more structured proteins due to relaxed selection on folding and a greater tolerance of mutational input^91^. At the same time, evolutionary conservation of structural disorder under stabilizing selection for the biophysical consequences of disorder can coexist with pronounced sequence divergence^55,92,93^. We did not find greater or more variable amino acid sequence divergence in more disordered proteins (Supplementary Fig. S12 ab). Yet, a greater binding plasticity of these proteins, that places them in a wider range of selective environments and promotes an accumulation of neutral variation under inconsistent or weak selection, covaried with their greater and more variable sequence divergence (Fig. 5 c, d). This corroborates the finding of greater binding plasticity in disordered proteins both within and across species (Figs. 3; 6a). On the technical side, the discordance between amino acid sequence and function in the ordered and disordered regions of a protein calls for a more nuanced approach to the evaluation of divergence of these proteins than multiple sequence alignment^29^. On the biological side, it provides a powerful mechanism of dimension reduction in developmental processes^51^, evidenced in the ability of IDPs to transduce processes operating at the level of cells to the scale of morphogenesis. Patterns operating in a relatively small population of cells early in development, such as studied here, can be influential in delineating morphogenesis in subsequent developmental expansion^59,94,95^. Together with path-resetting links in the core regulatory network (Fig. 3h), this contributes to the much sought-after internal consistency of mechanisms and rules across levels of developmental organization – from uniform cells to specialized structures.

## Methods

### Data collection and sample sizes

We measured cell morphology and protein expression in mesenchyme cells of upper beaks of *n* = 402 embryos across HH25-36 developmental stages and 22 study populations grouped into three genetically distinct groups (Supplementary Table S1). Protocols for incubation to the required developmental stage and storage of samples is in ^59^. Briefly, beaks were cryosectioned at 8 µm (approximately one-cell-thickness) and stored at -80°C. Thirteen sections per individual were obtained at beak midline: one section was stained with Alcian blue hematoxylin and eosin (H&E; U. Rochester MC) to delineate the histological area of interest, twelve were used in immunohistochemical (IHC) assays described below.

### Cell measurements and method validation

Cell measurements and protein expression data were collected within Area of Interest (AOI) of the upper beak that was delineated based on landmarks homologous across developmental stages and confirmed with H&E histological staining as described in ^65^. Using software we developed for this study^65^, which also see for tests of repeatability and validation of cell morphology and protein expression, a scalable 10x10 grid was overlayed on the AOI of each section and morphology was measured for every cell within each grid box. For each cell we recorded size (μm^2^), centroid coordinates, perimeter (μm), major and minor axes, length of the best fitting ellipse, aspect ratio, and feret length, angle and coordinates. We then assessed histological content of all 39,401 grid boxes containing 2,792,362 cells and retained only grid boxes fully comprised of undifferentiated mesenchymal cells. Grid boxes containing condensations, differentiating bone and cartilage tissues (at later stages), capillaries, epithelial boundaries, as well as boxes containing tissue gaps and folds were excluded from this study. Within the retained grid boxes, we further excluded cells that were overlapping, dividing, or damaged during sectioning, resulting in complete measurement and protein expression data for 724,465 mesenchymal cells in 5,886 grid boxes, each containing 50-330 cells (Supplementary Data S1, Supplementary Table S1).

### Classification of cell jamming/unjamming state

We classified cell jamming for each cell group (grid box) by calculating agreement among five metrics indicative of jamming transitions (Supplementary Fig. S1). Justification for using this method to infer cell jamming in 3D tissues is in^59^. Briefly, a combination of cell shape index (s.d. and mean) and cell aspect ratio (s.d. and mean) showed the closest agreement in identification of both jammed and unjammed states in multiple correspondence analysis (Supplementary Fig. S1). First, we calculated the cell shape index 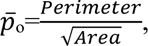 and used *p*\*=3.81 as a critical value below which cells are jammed, such that any cell rearrangements require changes in cell shape ^96^. Second, to examine the effect of shape elongation, we calculated cell aspect ratio 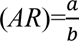 and its variability (standard derivation, s.d.) and used s.d. (*AR*) =0.5 (corresponding to *AR*≈1.6 in our sample) as a critical value below which cells are increasingly uniform and round, and are jammed^97^.

### Immunohistochemistry

For immunostaining, we used anti-β-catenin (610153, 1:16,000, BD Transduction Laboratories), anti-CaM (sc-137079, 1:15, Santa Cruz Biotechnology), anti-Wnt4 (ab91226, 1:800; Abcam), anti-Tgfβ2 (ab36495, 1:800, Abcam), anti-Bmp4 (ab118867, 1:100, Abcam), anti-Ihh (ab184624, 1:100, Abcam), anti-Dkk3 (ab214360, 1:100, Abcam), and anti-Fgf8 (89550, 1:50, Abcam) antibodies using methods described previously^59,65^. Validations showing specificity of stains are in^59^. Reactions were visualized with either diaminobenzidine (DAB, Elite ABC HRP Kit, PK-6100, Vector Labs) or Vector Red Alkaline Phosphatase substrate and Vectastain ABC-AP Kit (AK-5000, Vector Labs) and nuclei were counterstained with hematoxylin. Three slides, each containing four tissue sections (12 sections per embryo) were run with the following grouped antibodies: i) β-catenin, Fgf8, Tgfβ2 and no primary control, ii) Bmp4, Wnt4, Ihh and no primary control, and iii) Dkk3 and no primary control, and CaM and no primary control (Appendix S2). These were imaged and named according to embryo ID, protein, developmental stage, and IHC run to enable high-throughput processing as described in^65^. Across IHC runs, we randomized the assignment of sections from different populations and stages.

### Protein sequencing, multiple sequence alignments, and protein disorder analyses

The sequences for 20 genomes of house finches the study populations (Supplementary Table S1) were aligned to the *cibio_Scana_2019* version of the canary genome with BWA, and PCR duplicate reads were identified with Picard. Variant calls were made with GATK’s HaplotypeCaller and annotated with ANNOVAR using a database made from the canary assembly with its predicted transcripts. The resulting table was filtered for protein-altering exonic variants, including substitutions, splice site variants, and indels. We manually reviewed all these variants identified in the nine genes (Calm2 was included in addition to Calm1= CaM in IHC assays above) under this study by viewing the assembly in IGV. The variants that passed the review were then input into a custom script to make the corresponding substitutions specified by ANNOVAR in the predicted protein sequences.

Accession numbers for the nine genes were obtained from the Gene database on the NCBI portal. 170 bird species had records for one or more of these genes on 23 Feb 2023 (Supplementary Data S3-S11). The gene identifiers were then used in a custom script to fetch all records from the Gene database for each accession number in xml form and extract the transcript and protein accession numbers, which were filtered to retain only RefSeq curated (’NM_’ and ‘NP_’) and predicted (’XM_’ and ‘XP_’) sequences. The filtered accession numbers were then used to retrieve the associated amino acid sequences in fasta format from the Nucleotide database.

We aligned multiple sequences of each of nine proteins across species with Clustal Omega (*beta* 2023) using default settings. Percent identity matrices for each protein are shown in Supplementary Table S8. Evolutionary divergence was assessed based on *n* = 57 species with the full set of all nine proteins of comparable quality and is given in Supplementary Tables S8-9.

Binding affinity was assessed with STRING^98^. Intrinsic disorder for each protein sequence was visualized with AlphaFold3^99^ and agreement among five commonly used per-residue disorder predictors (PONDS VSL2, PONDR VXT, PONDR VL3, PONDR Fit, and IUPred) was calculated^100-107^ and given in Supplementary Figs. S3-11. Functional disorder analysis with positional information was conducted on the d2p2 platforms with output from nine predictors^108^. Propensity to form liquid-liquid phase separation was calculated with PSPredictor platform^109^. Data for focal proteins in all analyzed species are in Supplementary Table S7.

### Statistical analyses

To achieve normal distribution, reduce skewness and stabilize variance, we used the Box-Cox transformation with λ=0.5 for raw data on protein expression, log10 transformation for cell morphology measures and arcsine transformation for cell density proportional measures. We used a mixed-effects model to examine the effects of cell morphology or jamming state on protein expression. Population and embryo ID were treated as random effects. We computed the least-squared means of protein expression for cell mechanical states and assessed significance of these means and difference between them with a Sidak adjustment for multiple comparisons (Supplementary Table S4). To directly compare the contribution of both random and fixed effects to variance in protein expression (Supplementary Tables S2bc, S3b), we compared their contribution to Adjusted R². We first constricted a full model incorporating all fixed effects and then generated an array of reduced models, starting with the intercept-only model to assess baseline variance and sequentially omitting each predictor in other models. For each model, we extracted the residual variance estimates, and Adjusted R² was computed as: Adj R² = 1 - (Residual Variance (Reduced Model) / Residual Variance (Null Model)). The contribution of each predictor was then calculated as the difference in Adjusted R² between the full model and each reduced model and converted into percentages. This approach allowed us to quantify the independent explanatory power of each predictor while accounting for shared variance in the model. We then ranked these contributions to Adj R² for each predictor for Fig. 2 (Supplementary Tables S2-4). We derived matrices of pairwise partial correlations for each cell jamming state (Supplementary Table S5a, b) and divergence matrix between the cell states (Supplementary Table S5c) as well as matrices for each region (Supplementary Table S6-c) and pairwise and total divergence matrices among regions (Supplementary Table S6 d-g). Lack of dosage-dependence in binding plasticity of ordered proteins, and associated differences in linearity of correlations between ordered and disordered proteins, precludes the direct comparisons of eigenstructure of cell-state divergence and region-divergence matrices in Fig. 3. We thus calculated direct contribution of each element to matrix divergence between cell states and pairwise regions comparisons and overall divergence matrices as shown in Supplementary Tables S6h). We then ranked these contributions for each element for plotting on Fig. 3h.

## Data availability

All data are available in the manuscript and the supplementary materials.

## Author contributions

AVB designed the project and obtained funding; AVB, CAL, CMS, SEB, LJM, RAD developed methods, collected data, and analyzed data. AVB wrote original draft, AVB, CAL, CSM, SEB, RAD reviewed and edited final version.

## Supplementary Materials

### Supplementary text

#### Core network of transcriptional regulators in avian beak ontogeny

We examined the coexpression of nine key factors whose interactions play a central role in avian beak development: β-catenin, bone morphogenic protein 4 (Bmp4), calmodulin1 (Calm1), calmodulin2 (Calm2), dickkopf homolog 3 (Dkk3), fibroblast growth factor 8 (Fgf8), Indian hedgehog (Ihh), transforming growth factor beta 2 (TGFβII), and wingless type 4 (Wnt4). Below we review what is currently known about functional links and expression associations between these proteins. Figure 1a summarizes these findings.

Several conserved cellular signaling pathways converge on the regulation of downstream transcription factors specific to bone and cartilage differentiation and proliferation including osterix (OSX), runt-related transcription factor 2 (RUNX-2), and SRY-box transcription factor 9 (Sox9)^1^. OSX is an osteoblast specific transcription factor which works by committing cells to an osteoblast lineage^2^. RUNX-2 is a master regulator of osteoblast differentiation and targets genes specific to osteoblast differentiation and maturation^3^. Additionally, RUNX-2 is an important player in chondrocyte hypertrophy in endochondral bone formation (when bone replaces a cartilage template). Sox9 works by committing cells to a cartilage lineage and by transcribing genes necessary for the maturation of chondrocytes ^4^. Pathways that are known to regulate these genes include the canonical WNT signaling pathway^5-7^ as well as non-canonical WNT pathways, such as the Ca2+ pathway^8^. Additional regulatory pathways important in these tissues include BMP/TGFβ signaling^9-11^, mechanotransduction pathways^12^, and mitogen-activated protein kinase (MAPK) pathways^13^. Below we describe how the eight proteins investigated in this study interact in these pathways.

Wnt4 and β-catenin are key players in the canonical WNT signaling pathway^14^. When canonical WNTs are present and bind to cell surface receptors, cytoplasmic β-catenin is stabilized and as levels increase it is translocated to the nucleus and acts as a transcription factor targeting WNT genes^15^. Ihh may be an upstream activator of the canonical WNT signaling pathway, as shown in the osteogenesis of mouse endochondral bones^6^. The interaction between Ihh and the canonical WNT pathway may be tissue specific: in the context of osteoblast differentiation the WNT/β-catenin pathway is a downstream target of Ihh signaling while in chondrocytes, WNT/β-catenin is an upstream regulator of Ihh^16^. Ihh is integral to the formation of endochondral bones via its regulation of chondrocyte differentiation, proliferation, and maturation into osteoblasts^17,18^. Fgf8 is also a positive downstream target of the canonical WNT pathway, specifically in the upper jaw and rostrum during facial development in mice^19^.

On the other hand, DKKs are known WNT inhibitors^20-22^. Although Dkk3 is a divergent member of the family, it also likely inhibits WNTs^23^, and the canonical pathway specifically by inhibiting β-catenin attenuation^24,25^. In fact, Dkk3 may be a critical player in switching between the canonical and non-canonical WNT pathways. At the same time, Dkk3 can also be downstream of the canonical WNT signaling pathway^26^.

In the non-canonical Ca_2_+ pathway, Wnt4 works with CaM to activate the downstream calmodulin-dependent protein kinase II (CAMKII), which then activates NF-κB, a transcription factor involved in cell proliferation and differentiation^27^. The NF-κB pathway is specifically involved in differentiation and maturation of bone cells as well as chondrocyte differentiation^28^. Ihh can be downstream of the Ca_2_+ pathway, where it is a key determinant of chondrocyte hypertrophy^29^. Fgf8 can also activate NF-κB, which then upregulates gelatinases in the cartilage and associated tissue remodeling^30^.

The canonical and non-canonical WNT pathways can antagonize and regulate one another. In chicks, CAMKII is active in proliferating chondrocytes before they become hypertrophic but negative regulation from the canonical pathway allows for the work of genes like *RUNX2,* leading to chondrocyte hypertrophy^29^. On the other hand, CAMKII can regulate the canonical pathway in part via phosphorylation and suppression of β-catenin^31,32^.

MAPK pathways are another set of cellular signaling systems that have been implicated in bone and cartilage development. Wnt4 can activate the p38 MAPK pathway by directly activating p38, inducing a cascade which activates genes involved in bone regeneration^33^. Other factors can also act via MAPK signaling in craniofacial development including Fgf8 and TGFβII ^34,35^. For example, Fgf8 regulates chondrogenesis in the chick mandible specifically via MEK-ERK activation of the MAPK cascade^36^. Downstream targets of the p38 and ERK MAPK pathways include *RUNX2* and *OSX* ^34^.

Another major set of pathways involved in bone and cartilage development are the BMP/TGFβ signaling pathways^9^. BMPs and TGFβs both have canonical pathways in which ligands bind to cell membrane receptors and initiate the phosphorylation of SMAD proteins, ultimately resulting in a complex that translocates to the nucleus and induces transcription for bone and cartilage specific genes^1,37^. Two of our studied proteins, Bmp4 and TGFβII, operate in this pathway. Dkk3 may enhance TGFβ signaling in chondrocytes, as seen in human cell cultures^22^. Dkk3 can also be downstream of BMP/TGFβ signaling pathways^26^.

The BMP/TGFβ pathway and canonical WNT pathway can either work together or antagonize each other, which may be stage- or tissue-dependent ^38^. For example, in chondrogenesis, TGFβ signaling enhances WNT signaling in immature chondrocytes but antagonizes signaling in mature chondrocytes ^39^. One mechanism by which these pathways can cooperate occurs when TGFβII activates β-catenin signaling in the canonical pathway via the action of Smad-3 and Smad-4, leading to expression of chondrocyte specific genes^37^. In osteogenesis, the pathways work together for osteoblastic differentiation, but once in osteocytes, these pathways play opposite roles in regulating osteoclast genesis and bone resorption^38^. Specifically, Bmp4 is downstream of WNT/ β-catenin signaling while still in the osteoblast precursor stage in mice^40^. Similarly, multiple WNT family members are known to activate BMP signaling in zebrafish craniofacial development^41^.

Ihh and Fgf8 also have important interactions with BMP signaling. In some contexts Ihh is needed for BMP induced osteogenesis^42,43^. In mice, Ihh knockouts led to reduced Bmp4 expression in the skull^44^ and in chicks Ihh upregulates Bmp4, leading to chondrocyte differentiation^45^. Bmp4 and Fgf8 can inhibit each other^46,47^. In the mandibular processes of developing mice Fgf8 suppresses the expression of Bmp4 via the transcription factor PITX2^46^. Bmp4 in the ventral part of the head in chicks can limit Fgf8 expression^48^. By acting antagonistically these two factors have their own expression domains which may be important for patterning very early in craniofacial development^48^. Many FGF family members are also known to be Ihh inhibitors^16^, leading to complex interactions between these factors.

Many of our factors of interest have been implicated in avian beak development specifically. Bmp4 has many known associations with beak size and shape^49,50^. For example, in Darwin’s finches increased Bmp4 expression is associated with wider and deeper beaks and larger beak sizes are associated with increased expression domains of β-catenin, Dkk3, and the TGFβII receptor^26,51^. Additionally, CaM expression is associated with longer, pointed beaks, whereas decreased expression is associated with shorter, more robust beaks^52^, and Fgf8 in the epithelium is necessary for elongation of the beak in early to mid-developmental stages via increased chondrogenesis^53^. In Caribbean bullfinches, Ihh and Bmp4 expression are correlated with increased outgrowth of the premaxillary bone (pmx), and functional experiments in chickens confirm that increased expression of these factors increases the size of the pmx^54^. Further, Wnt4 is mainly expressed in the epithelium in the mid to early stages of chick craniofacial development^55^. These functional associations are summarized in Fig. 1a.

---

**Fig. S1.**
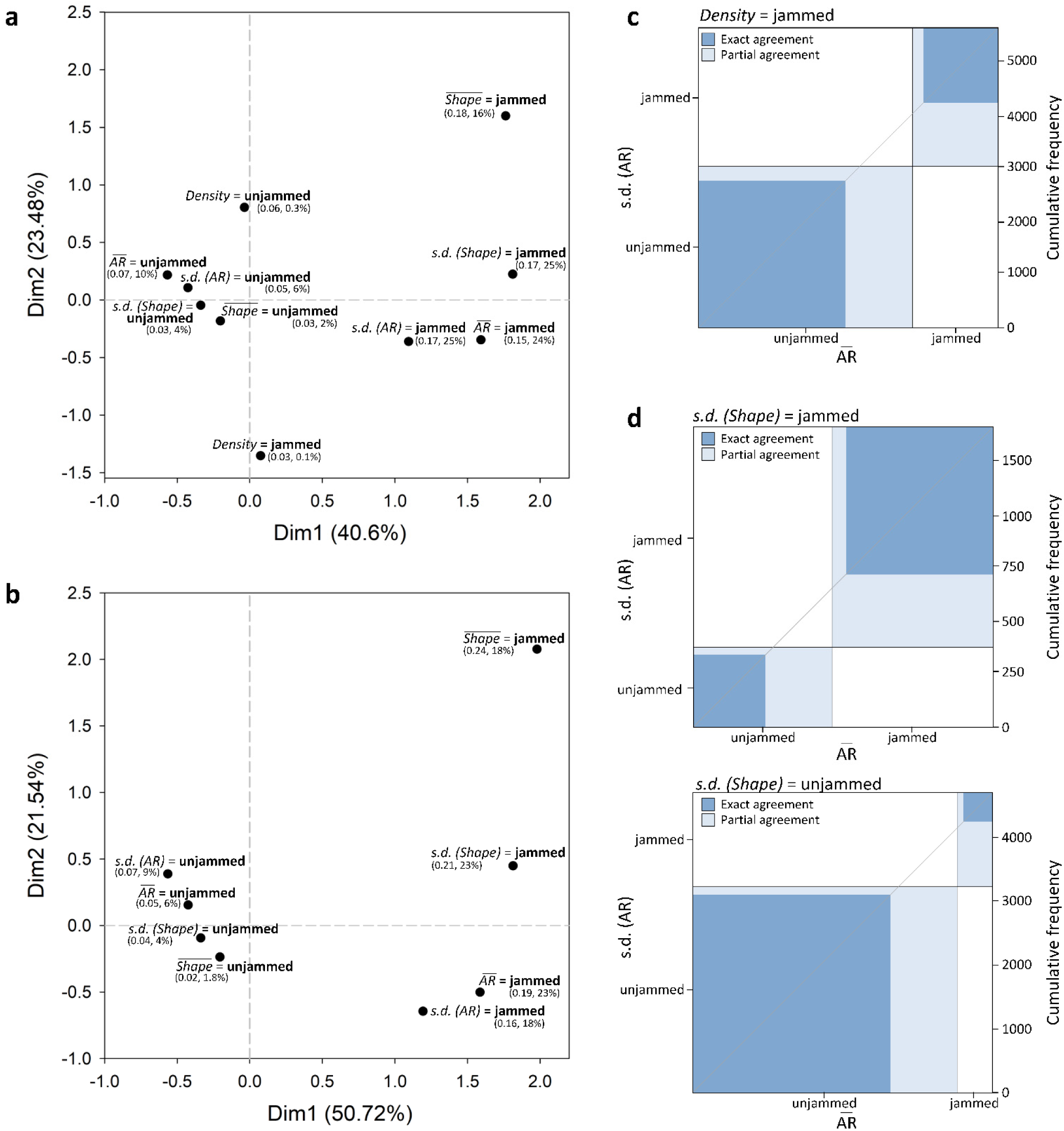
A combination of cell shape index (s.d. and mean) and cell aspect ratio (s.d. and mean) show the closest agreement in identification of both jammed and unjammed cell states in multiple correspondence analysis. **(a)** Cell density index does not distinguish dense migrating and dense jammed states (it contributes only to dimension 2) and thus agrees with other measures only in identifying unjammed state as shown in the Kappa plot **(c)**. **(b)** Cell shape and aspect ratio measures agree in identifying both jammed and unjammed states – they contribute equally to state separation among both dimensions as shown in the Kappa plots **(d)**. (a, b) Shown are variance accounted for by each dimension (axes) and partial contribution of each index (value and %) to the total chi-squared association among the indexes. s.d. (AR) shows better agreement with s.d. (Shape) in identifying cell state than mean (AR).

**Fig. S2.**
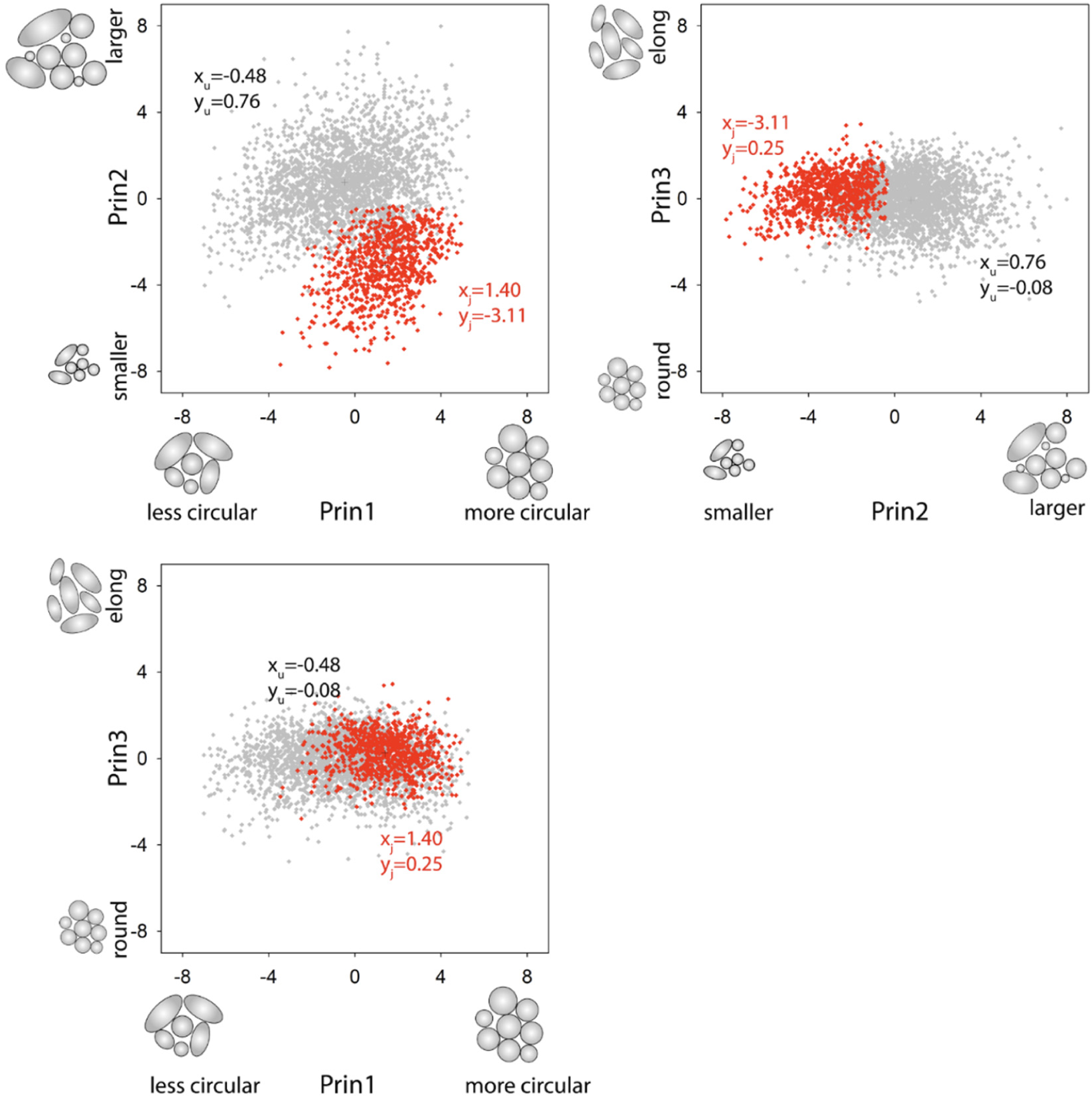
Cell groups classified as jammed (red, j) and unjammed (grey, u) according to the combination of cell shape index and AR (Fig. S1) consistently differ in morphology. Principal component analysis from Supplementary Table S2. Crosses show centroid coordinates for each group.

**Fig. S3.**
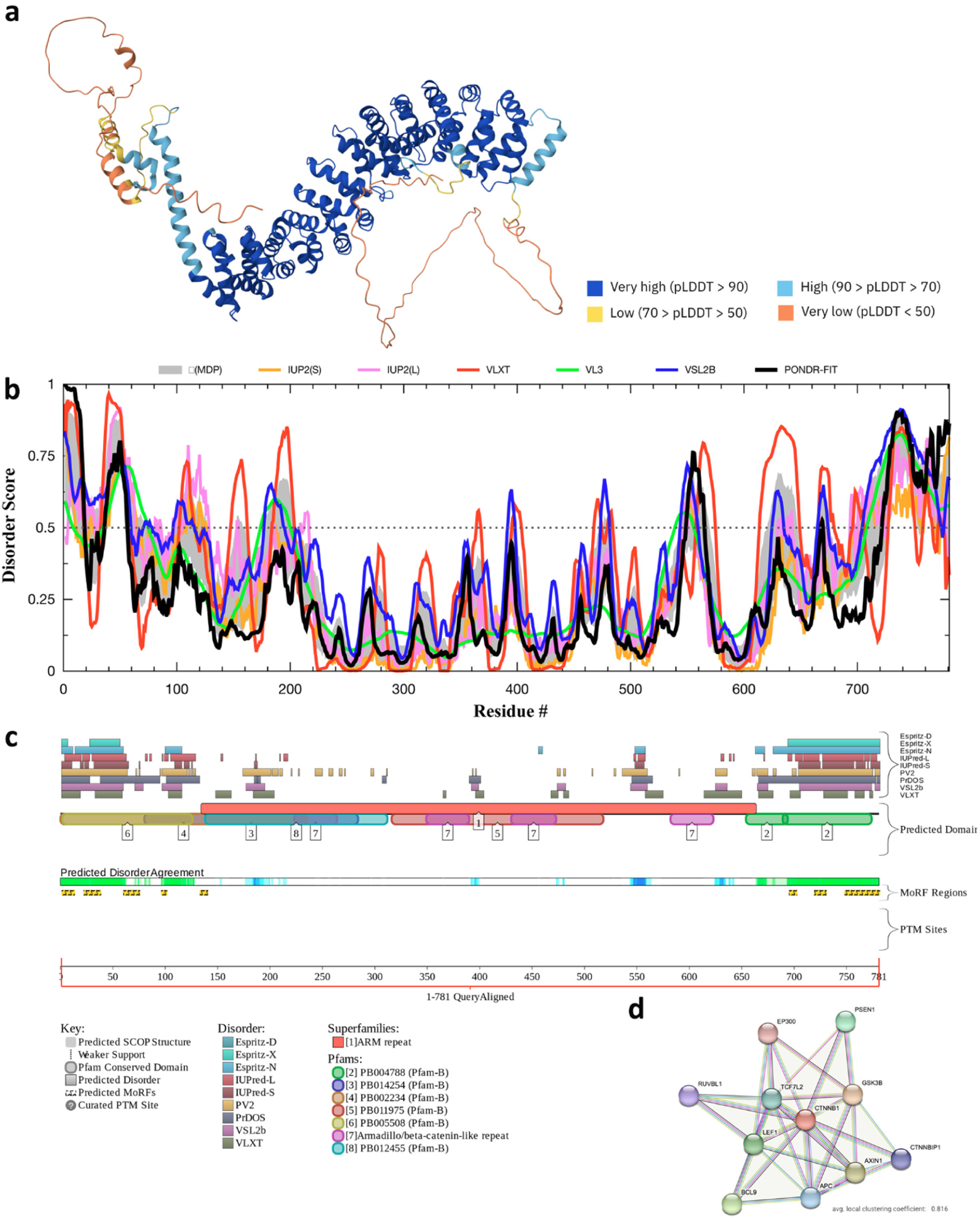
Evaluation of intrinsic disorder of the house finch CTNNB1 (b-catenin). **(a)** AlphaFold 3 model showing per-residue confidence scores (pLDDT). Regions with pLDDT < 50 are unstructured. **(b)** Disorder profile generated by RIDAO aggregating the results of MDP, UUP, VLXT, VL3, VSL2B, and PONDR-FIT per residue predictors of disorder (mean (line) ± s.e (shade); see Methods), regions <0.5 value for all predictors are disordered. **(c)** Functional disorder profile (d^2^p^2^ platform) based on the consensus of nine disorder predictors (green gradient). Blue gradient shows locations where the disorder predictions intersect the functional domain. Molecular recognition features (MoRF regions)- binding sites predicted to undergo disorder transition at binding with the specific partners are shown with yellow ZZZs (see Methods). **(d)** Binding affinity evaluated by STRING (see Methods) shows 33 edges, average node degree 6, and average local clustering coefficient = 0.816.

**Fig. S4.**
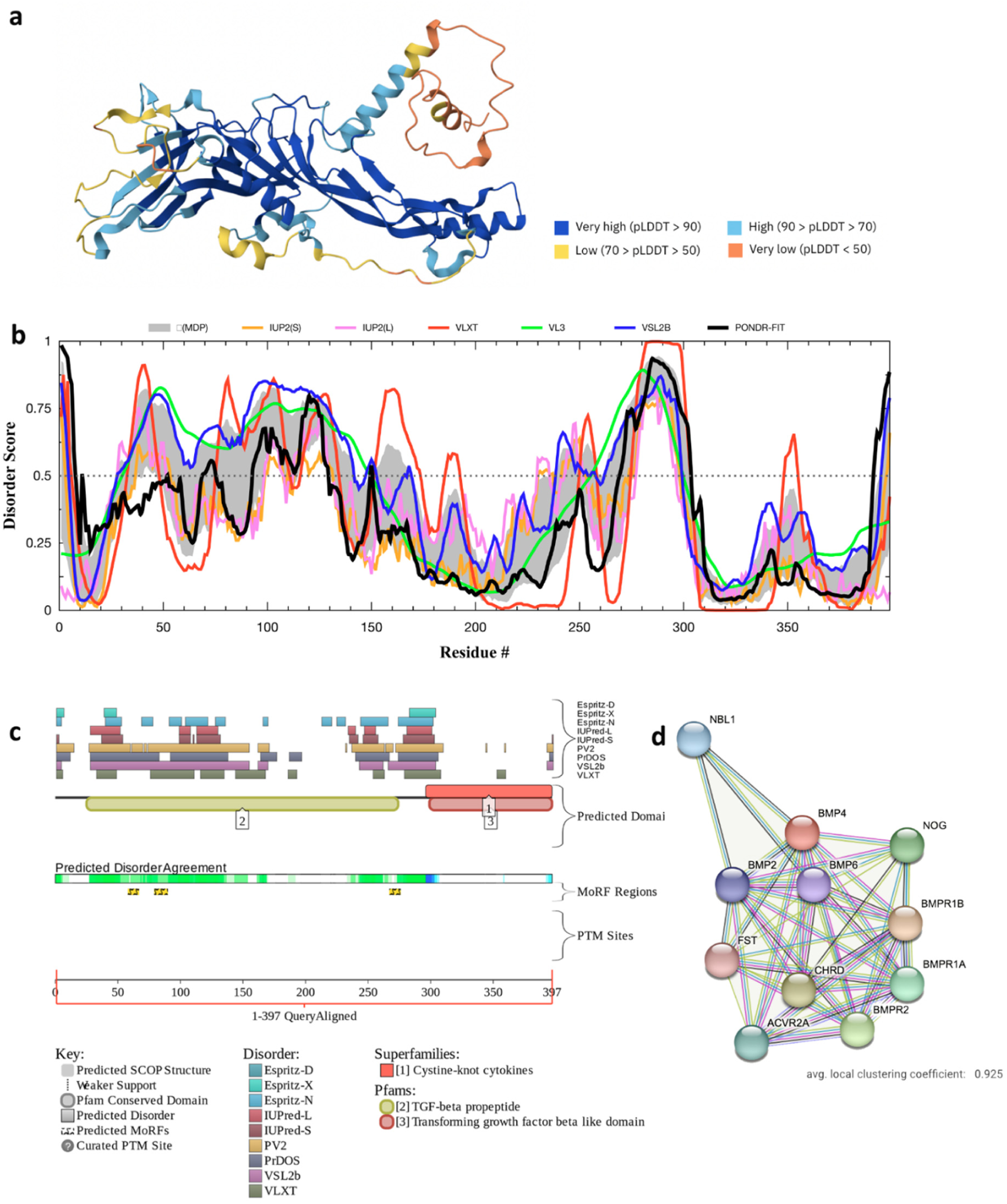
Evaluation of intrinsic disorder of the house finch Bmp4. **(a)** AlphaFold 3 model showing per-residue confidence scores (pLDDT). Regions with pLDDT < 50 are unstructured. **(b)** Disorder profile generated by RIDAO aggregating the results of MDP, UUP, VLXT, VL3, VSL2B, and PONDR-FIT per residue predictors of disorder (mean (line) ± s.e (shade)), regions <0.5 value for all predictors are disordered. **(c)** Functional disorder profile (d^2^p^2^ platform) based on the consensus of nine disorder predictors (green gradient). Blue gradient shows locations where the disorder predictions intersect the functional domain. Binding sites predicted to undergo disorder transition at binding with the specific partners are shown with yellow ZZZs. **(d)** Binding affinity evaluated by STRING shows 46 edges, average node degree 8.36, and average local clustering coefficient 0.925.

**Fig. S5.**
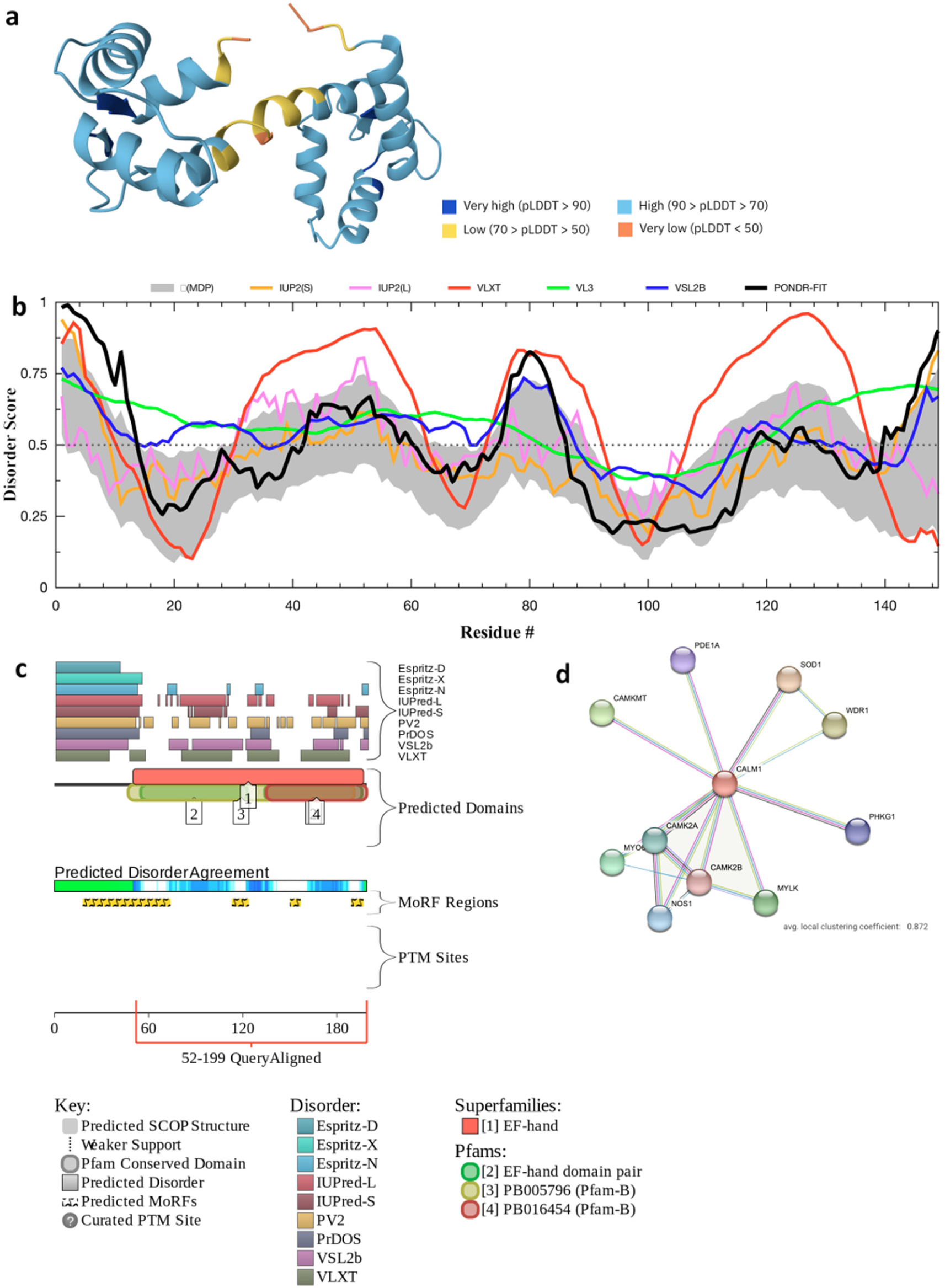
Evaluation of intrinsic disorder of the house finch Calm1 (calmodulin1). **(a)** AlphaFold 3 model showing per-residue confidence scores (pLDDT). Regions with pLDDT < 50 are unstructured. **(b)** Disorder profile generated by RIDAO aggregating the results of MDP, UUP, VLXT, VL3, VSL2B, and PONDR-FIT per residue predictors of disorder (mean (line) ± s.e (shade)), regions <0.5 value for all predictors are disordered. **(c)** Functional disorder profile (d2p2 platform) based on the consensus of nine disorder predictors (green gradient). Blue gradient shows locations where the disorder predictions intersect the functional domain. Binding sites predicted to undergo disorder transition at binding with the specific partners are shown with yellow ZZZs. (d) Binding affinity evaluated by STRING shows 17 edges, average node degree 3.09, and average local clustering coefficient 0.872.

**Fig. S6.**
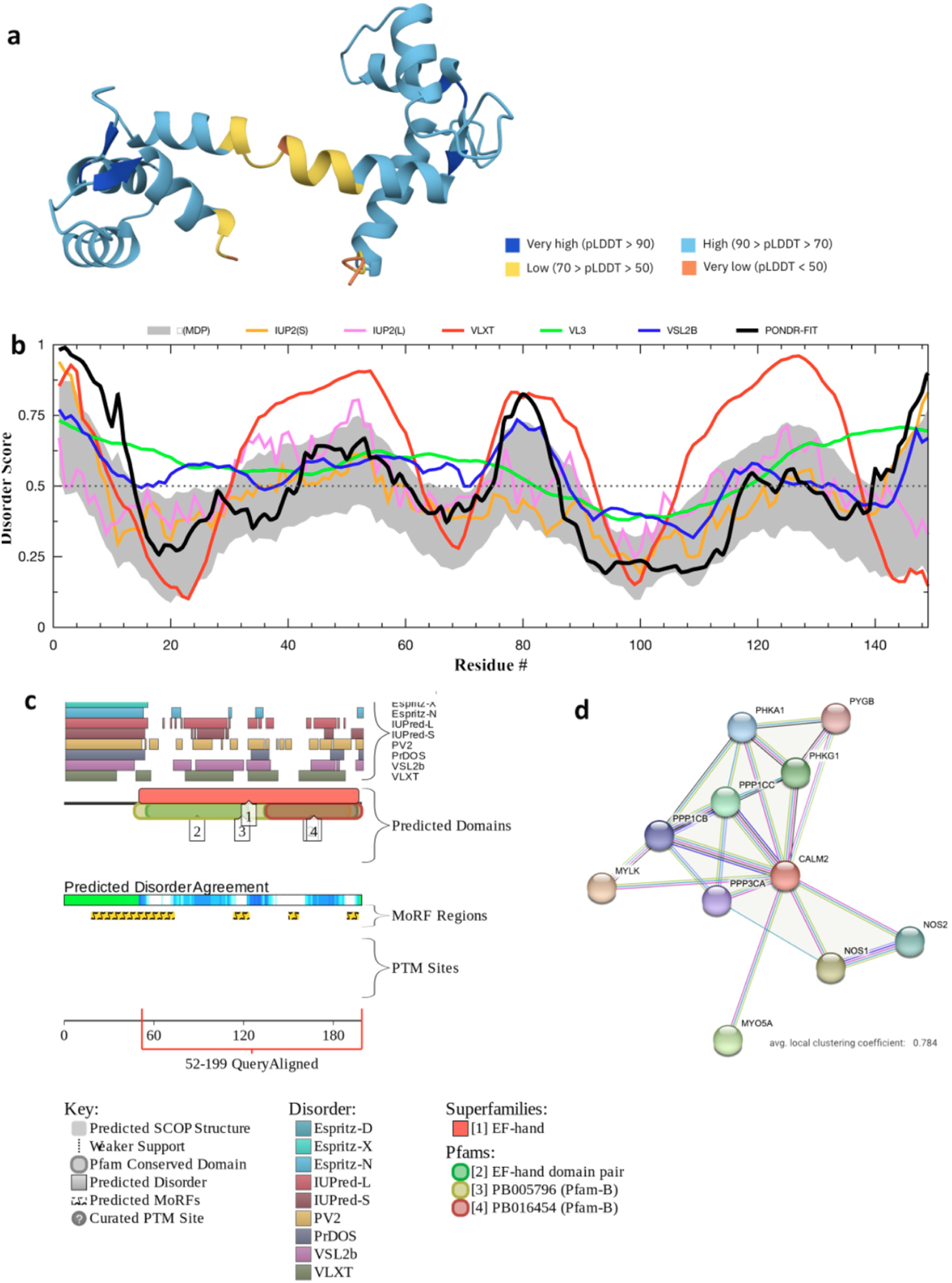
Evaluation of intrinsic disorder of the house finch CALM2 (calmodulin2). **(a)** AlphaFold 3 model showing per-residue confidence scores (pLDDT). Regions with pLDDT < 50 are unstructured. **(b)** Disorder profile generated by RIDAO aggregating the results of MDP, UUP, VLXT, VL3, VSL2B, and PONDR-FIT per residue predictors of disorder (mean (line) ± s.e (shade)), regions <0.5 value for all predictors are disordered. **(c)** Functional disorder profile (d2p2 platform) based on the consensus of nine disorder predictors (green gradient). Blue gradient shows locations where the disorder predictions intersect the functional domain. Binding sites predicted to undergo disorder transition at binding with the specific partners are shown with yellow ZZZs. (d) Binding affinity evaluated by STRING shows 23 edges, average node degree 4.13, and average local clustering coefficient 0.784.

**Fig. S7.**
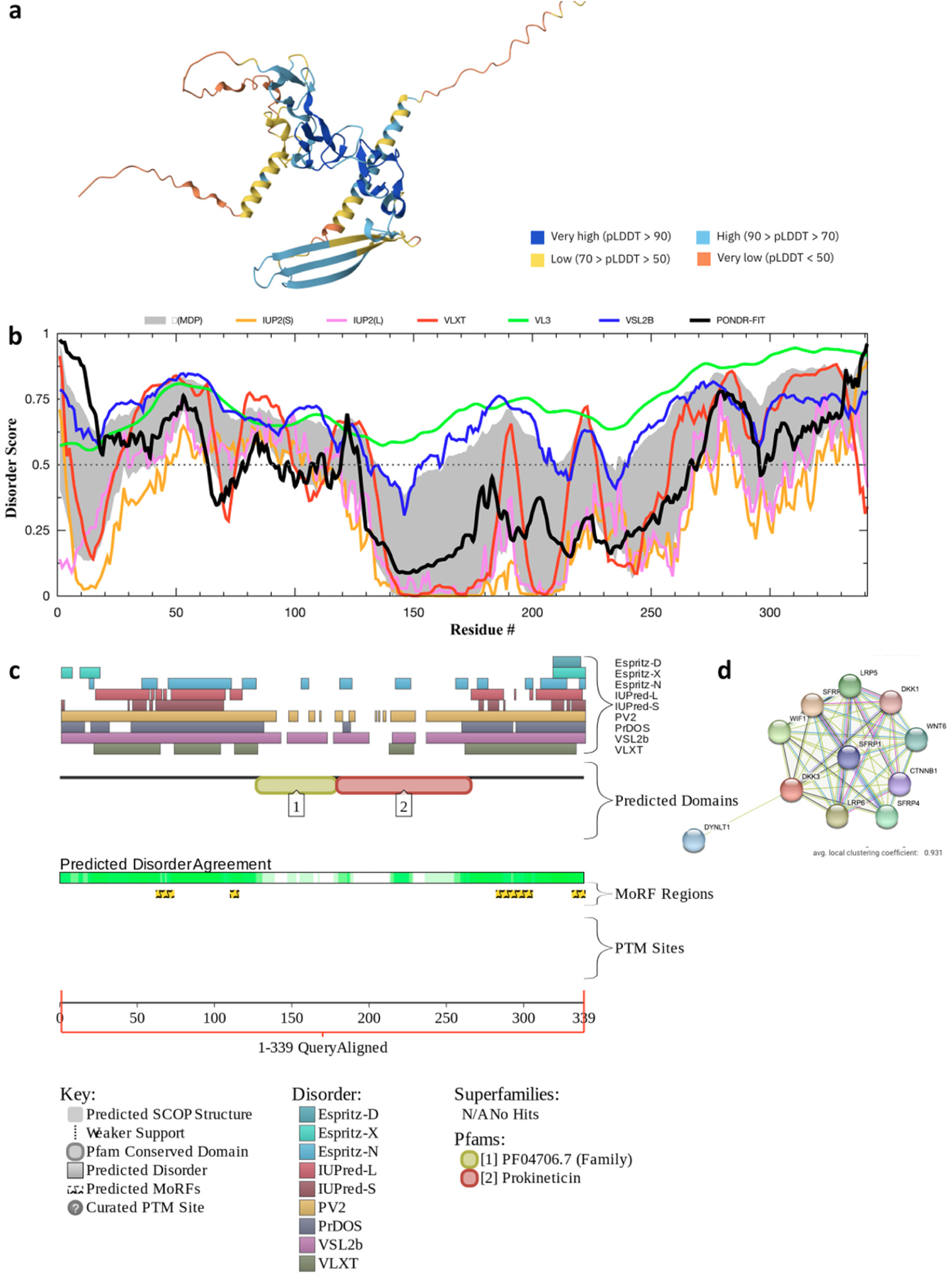
Evaluation of intrinsic disorder of the house finch Dkk3. **(a)** AlphaFold 3 model showing per-residue confidence scores (pLDDT). Regions with pLDDT < 50 are unstructured. **(b)** Disorder profile generated by RIDAO aggregating the results of MDP, UUP, VLXT, VL3, VSL2B, and PONDR-FIT per residue predictors of disorder (mean (line) ± s.e (shade)), regions <0.5 value for all predictors are disordered. **(c)** Functional disorder profile (d2p2 platform) based on the consensus of nine disorder predictors (green gradient). Blue gradient shows locations where the disorder predictions intersect the functional domain. Binding sites predicted to undergo disorder transition at binding with the specific partners are shown with yellow ZZZs. (d) Binding affinity evaluated by STRING shows 43 edges, average node degree 7.82, and average local clustering coefficient 0.931.

**Fig. S8.**
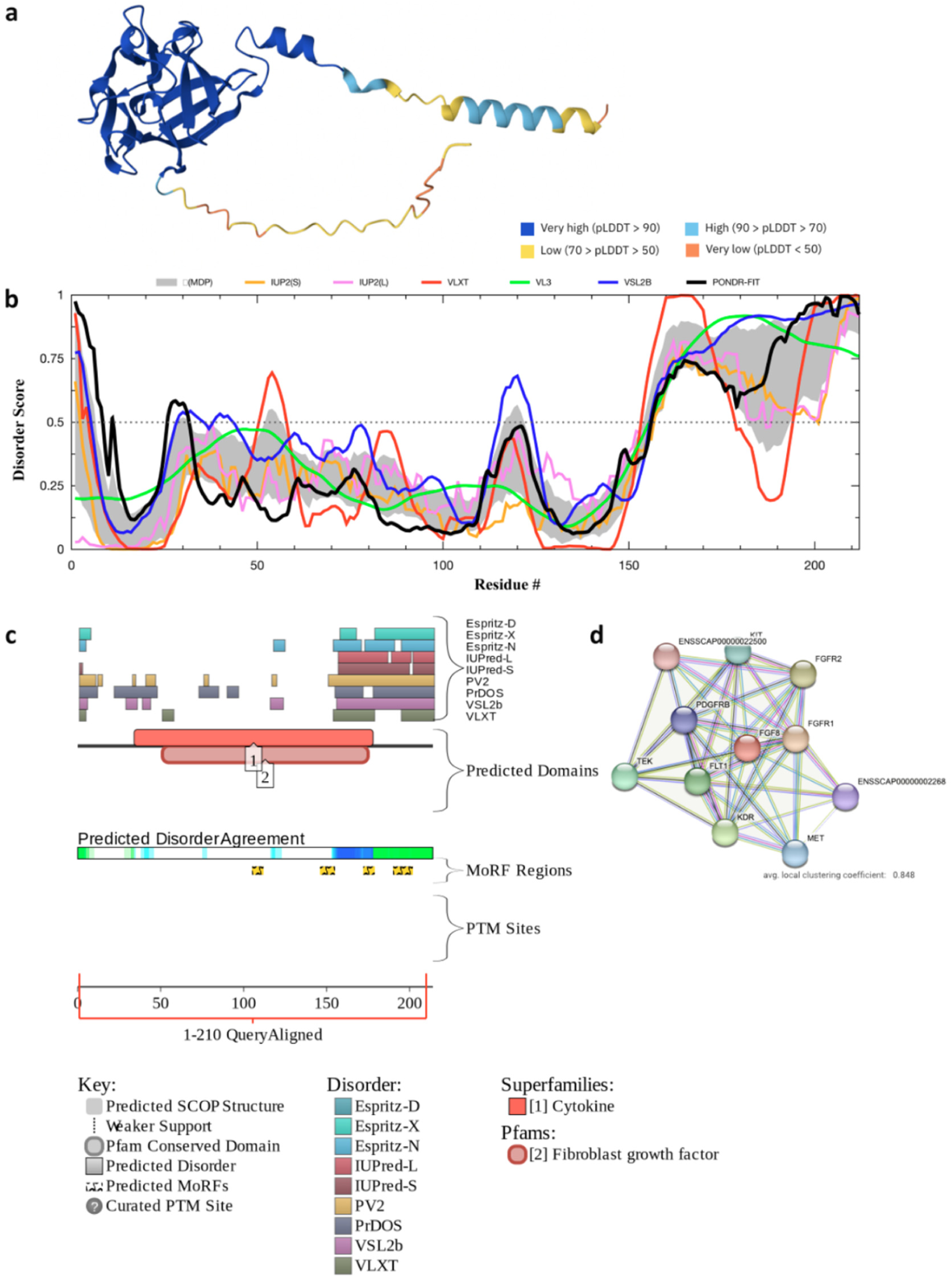
Evaluation of intrinsic disorder of the house finch Fgf8. **(a)** AlphaFold 3 model showing per-residue confidence scores (pLDDT). Regions with pLDDT < 50 are unstructured. **(b)** Disorder profile generated by RIDAO aggregating the results of MDP, UUP, VLXT, VL3, VSL2B, and PONDR-FIT per residue predictors of disorder (mean (line) ± s.e (shade)), regions <0.5 value for all predictors are disordered. **(c)** Functional disorder profile (d2p2 platform) based on the consensus of nine disorder predictors (green gradient). Blue gradient shows locations where the disorder predictions intersect the functional domain. Binding sites predicted to undergo disorder transition at binding with the specific partners are shown with yellow ZZZs. (d) Binding affinity evaluated by STRING shows 45 edges, average node degree 8.18, and average local clustering coefficient 0.848.

**Fig. S9.**
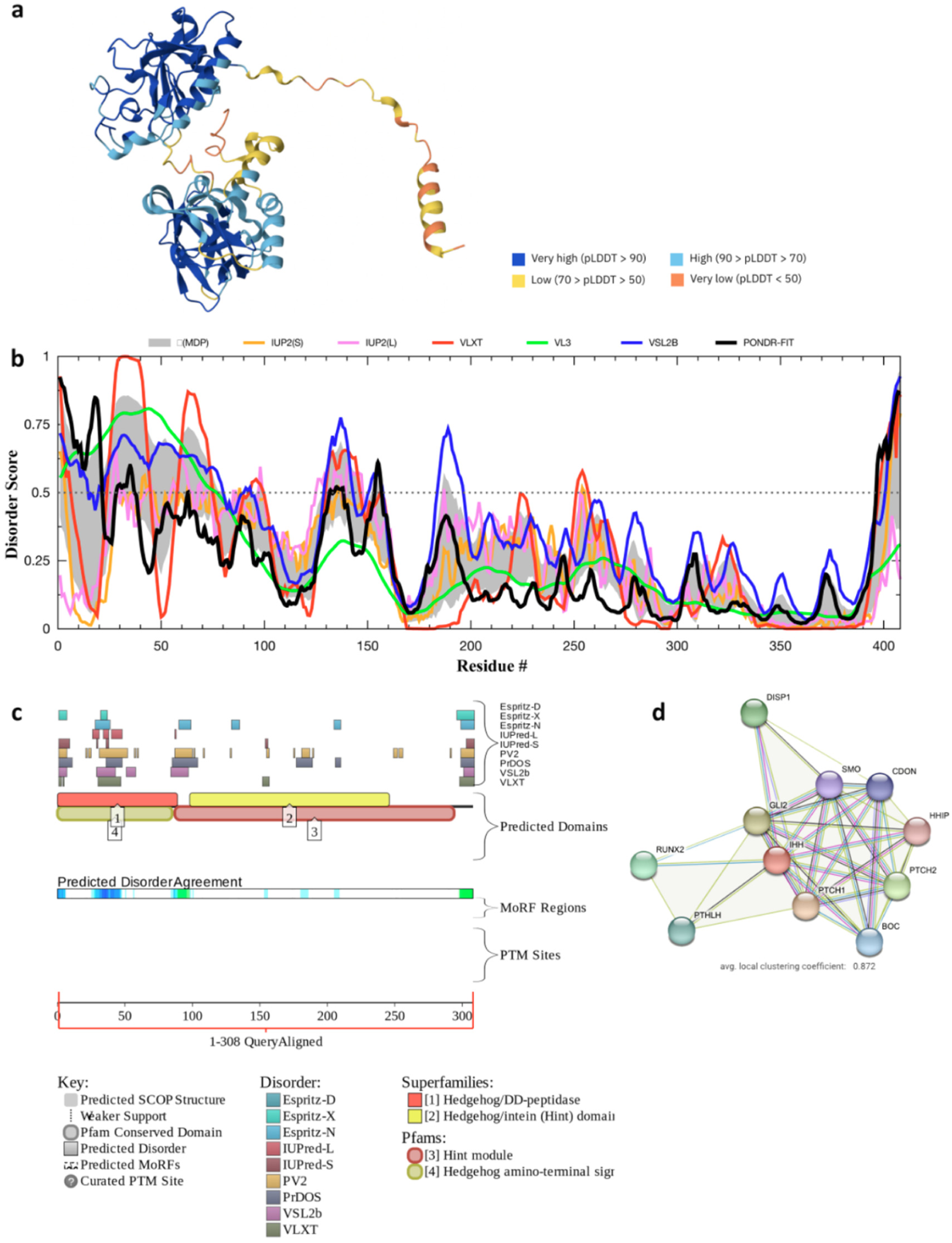
Evaluation of intrinsic disorder of the house finch Ihh. **(a)** AlphaFold 3 model showing per-residue confidence scores (pLDDT). Regions with pLDDT < 50 are unstructured. **(b)** Disorder profile generated by RIDAO aggregating the results of MDP, UUP, VLXT, VL3, VSL2B, and PONDR-FIT per residue predictors of disorder (mean (line) ± s.e (shade)), regions <0.5 value for all predictors are disordered. **(c)** Functional disorder profile (d2p2 platform) based on the consensus of nine disorder predictors (green gradient). Blue gradient shows locations where the disorder predictions intersect the functional domain. Binding sites predicted to undergo disorder transition at binding with the specific partners are shown with yellow ZZZs. (d) Binding affinity evaluated by STRING shows 38 edges, average node degree 6.21, and average local clustering coefficient 0.872.

**Fig. S10.**
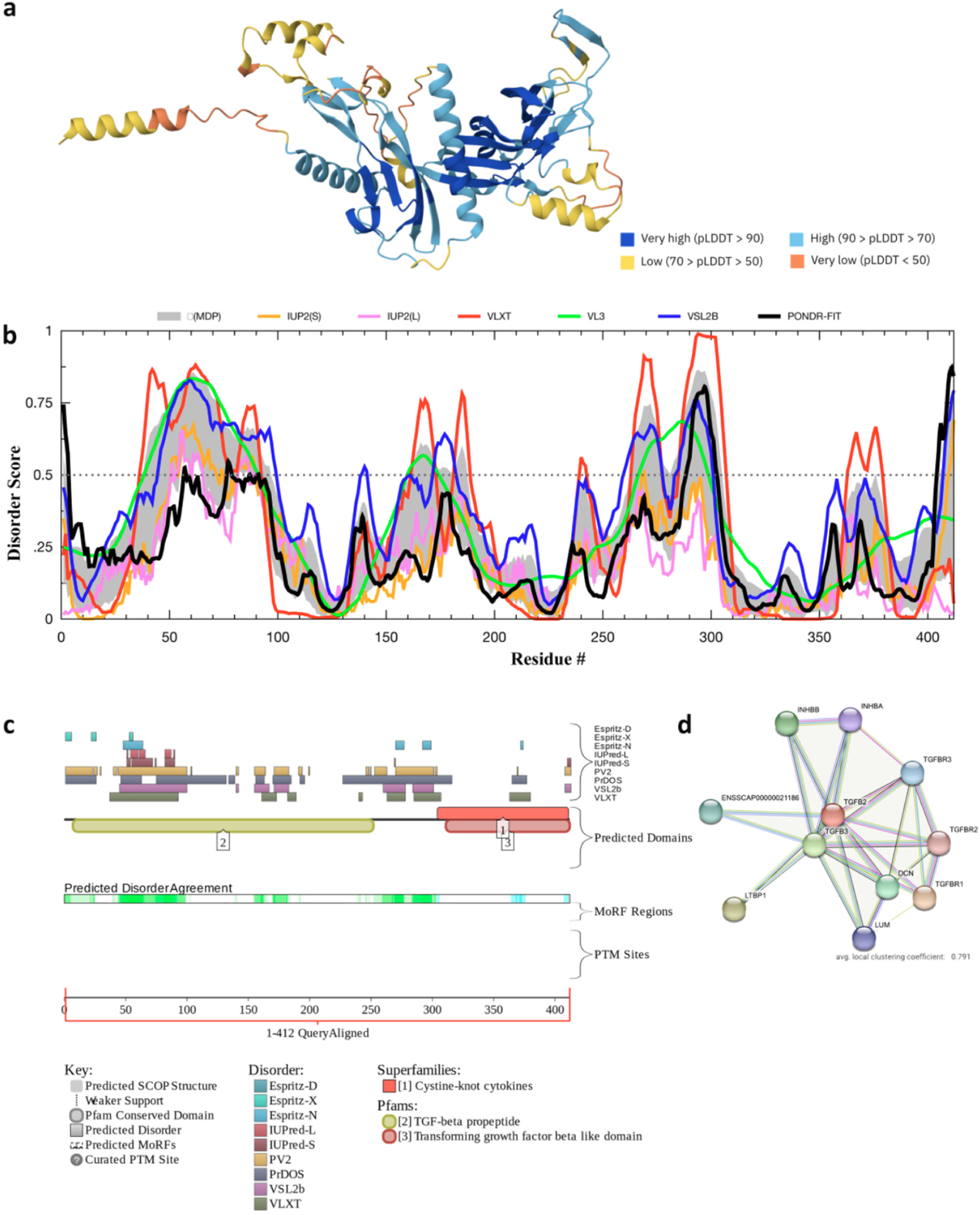
Evaluation of intrinsic disorder of the house finch TGFB2. **(a)** AlphaFold 3 model showing per-residue confidence scores (pLDDT). Regions with pLDDT < 50 are unstructured. **(b)** Disorder profile generated by RIDAO aggregating the results of MDP, UUP, VLXT, VL3, VSL2B, and PONDR-FIT per residue predictors of disorder (mean (line) ± s.e (shade)), regions <0.5 value for all predictors are disordered. **(c)** Functional disorder profile (d2p2 platform) based on the consensus of nine disorder predictors (green gradient). Blue gradient shows locations where the disorder predictions intersect the functional domain. Binding sites predicted to undergo disorder transition at binding with the specific partners are shown with yellow ZZZs. (d) Binding affinity evaluated by STRING shows 28 edges, average node degree 5.09, and average local clustering coefficient 0.791.

**Fig. S11.**
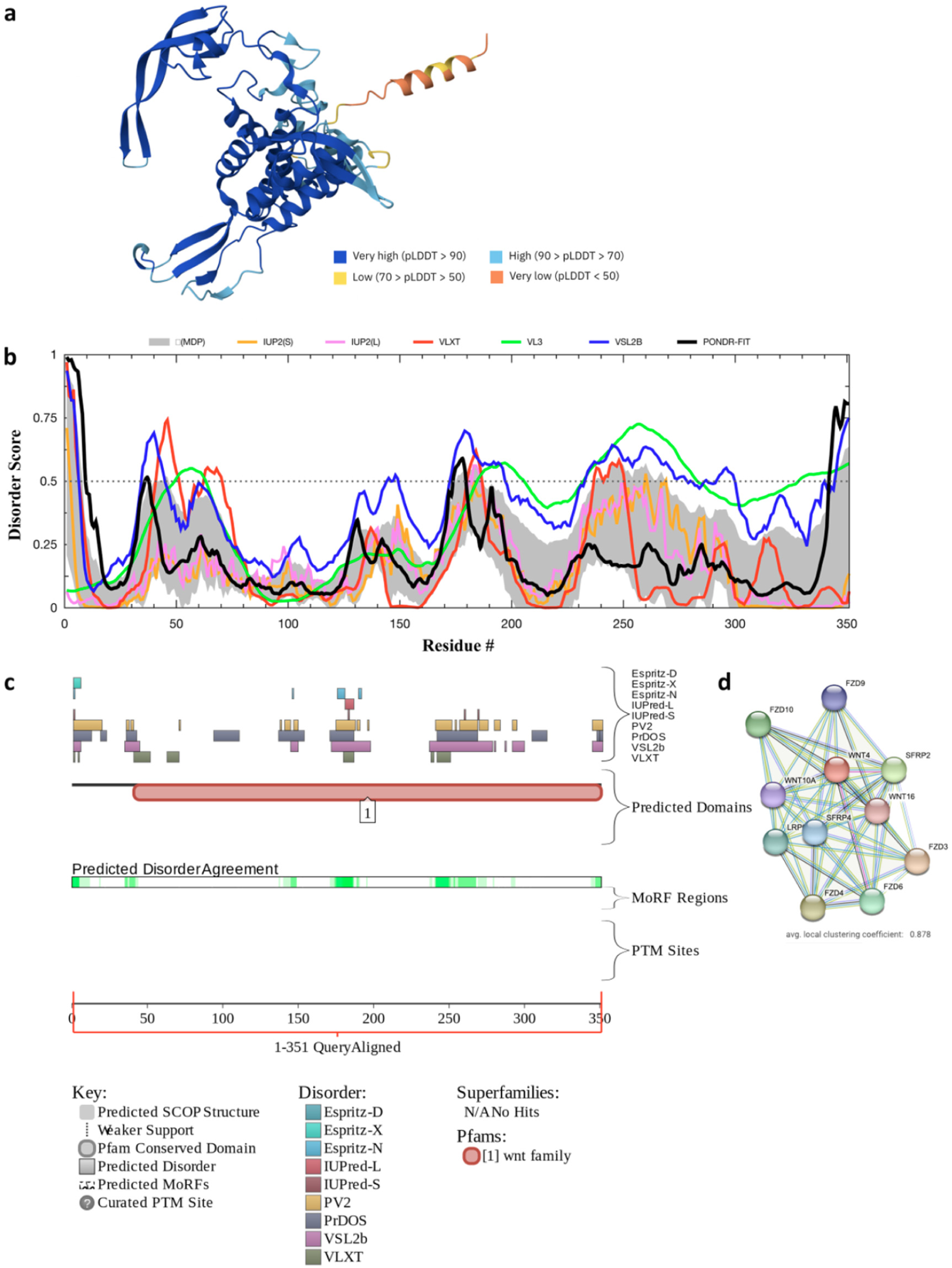
Evaluation of intrinsic disorder of the house finch Wnt4. **(a)** AlphaFold 3 model showing per-residue confidence scores (pLDDT). Regions with pLDDT < 50 are unstructured. **(b)** Disorder profile generated by RIDAO aggregating the results of MDP, UUP, VLXT, VL3, VSL2B, and PONDR-FIT per residue predictors of disorder (mean (line) ± s.e (shade)), regions <0.5 value for all predictors are disordered. **(c)** Functional disorder profile (d2p2 platform) based on the consensus of nine disorder predictors (green gradient). Blue gradient shows locations where the disorder predictions intersect the functional domain. Binding sites predicted to undergo disorder transition at binding with the specific partners are shown with yellow ZZZs. (d) Binding affinity evaluated by STRING shows 47 edges, average node degree 8.55, and average local clustering coefficient 0.878.

**Fig. S12.**
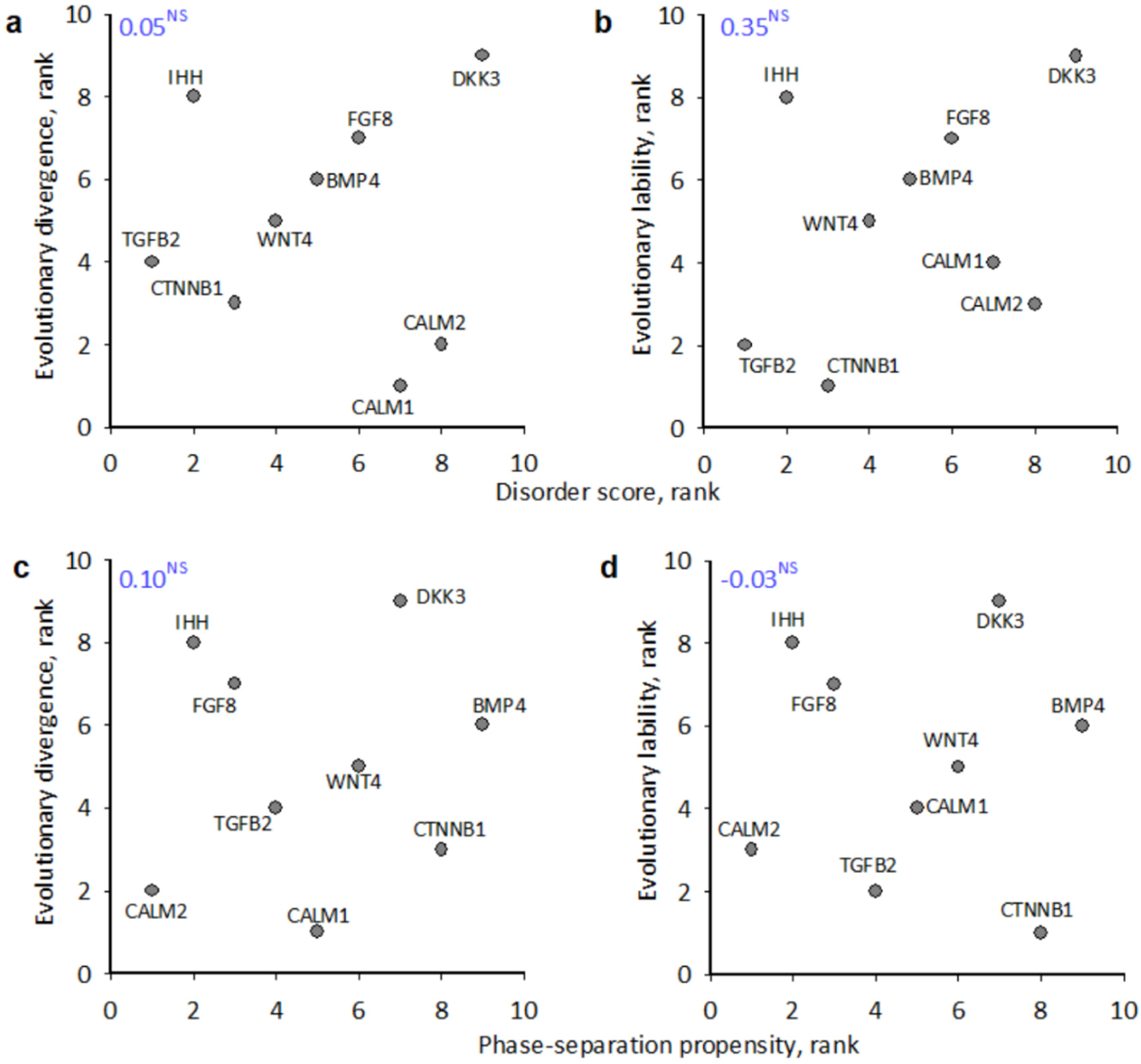
Neither disorder score (a, b) nor phase-separation propensity (c,d) are associated with evolutionary divergence of proteins under this study. Shown are Kendall’s correlation coefficients, NS designates lack of statistical significance. Data from Supplementary Tables S6-7.

**Supplementary Table S1.** Combined dataset of embryo samples.

| Region<br>**abbreviation**<br>*total embryos* | Sampling locations<br>latitude, °N, longitude, °W)<br>within a region | # *grid*<br>*boxes* * | Developmental stages (HH)<br>*embryos* |  |  |  |
| --- | --- | --- | --- | --- | --- | --- |
|  |  |  | 25-29 | 30-32 | 33-34 | 35-36 |
| Northern Montana<br>**nMT**<br>*n* = 134 | Fairfield (47.615545, 111.98016) | 42 | 10 | 24 | 12 | 18 |
|  | Great Falls (47.49839, 111.22444) | 655 |  |  |  |  |
|  | Cut Bank (48.629812, 112.333318) | 244 |  |  |  |  |
|  | Choteau (47.81437, 112.19193) | 15 |  |  |  |  |
|  | <u>Valier</u> (48.305658, 112.251628) | 206 |  |  |  |  |
|  | Lima (44.6369°, 112.5920°) | 66 |  |  |  |  |
|  | Havre (48.5500, 109.6841) | 20 | 12 | 28 | 24 | 6 |
|  | Malta (48.352662, 107.880582) | 1,295 |  |  |  |  |
| Eastern Montana<br>**eMT**<br>*n* = 145 | Plentywood (48.77368, 104.547787) | 17 | 9 | 83 | 37 | 16 |
|  | Glasgow (48.202811, 106.634599) | 1,129 |  |  |  |  |
|  | Circle (47.4156, 105.5907) | 314 |  |  |  |  |
|  | <u>Culbertson</u> (48.147585, 104.514955) | 694 |  |  |  |  |
|  | Poplar (48.1131°, 105.1983°) | 228 |  |  |  |  |
|  | Wolf Point (48.0906°, 105.6406°) | 107 |  |  |  |  |
|  | <u>Broadus</u> (45.4439°, 105.4075°) | . |  |  |  |  |
|  | <u>Billings</u> (45.7833°, 108.5007°) | . |  |  |  |  |
| Western Montana<br>**wMT**<br>*n* = 123 | <u>Kalispell</u> (48.21943, 114.33305) | 372 | 3 | 38 | 33 | 10 |
|  | Polson (47.687447, 114.160314) | 373 |  |  |  |  |
|  | Superior (47.193045, 114.888817) | 34 |  |  |  |  |
|  | Plains (47.459956, 114.884694) | 462 |  |  |  |  |
|  | Thompson Falls (47.600191, 115.342068) | 232 |  |  |  |  |
|  | <u>Missoula</u> (46.8787, 113.9966) | 590 | 5 | 16 | 18 | - |
|  | <u>Paws Up</u> (46.916957, 113.435576) | 157 |  |  |  |  |
|  | Hamilton (46.246863, 114.173587; | 70 |  |  |  |  |
| Total: |  |  |  |  |  |  |
| *n* = 352 embryos |  | *n* = 7,322 | 39 | 189 | 124 | 50 |

\*with pre-condensation mesenchymal cells only, underlined populations indicate sequenced genomes (*n* = 5 males per each populations)

**Supplementary Table S2a.** Principal component analysis of cell morphology based on *n* = 724,465 mesenchymal cells. Bold values show the largest projection on the original variable. Prin1 mostly captures variation in cell circularity, Prin2 – cell size, and Prin3 – cell elongation.

| Variable | Prin1<br>*circularity* | Prin2<br>*size* | Prin3<br>*elongation* |
| --- | --- | --- | --- |
| Area | 0.33 | **0.65** | 0.19 |
| Shape | **0.48** | -0.17 | -0.08 |
| Perimeter | 0.42 | **0.46** | 0.11 |
| Aspect Ratio (AR) | 0.32 | -0.51 | **0.72** |
| Circularity | **-0.48** | 0.13 | 0.07 |
| Solidity | -0.40 | 0.25 | **0.64** |
| Eigenvalue | 4.12 | 1.22 | 0.52 |
| Variance explained | 68.7% | 20.4% | 8.7% |

**Supplementary Table S2b.**
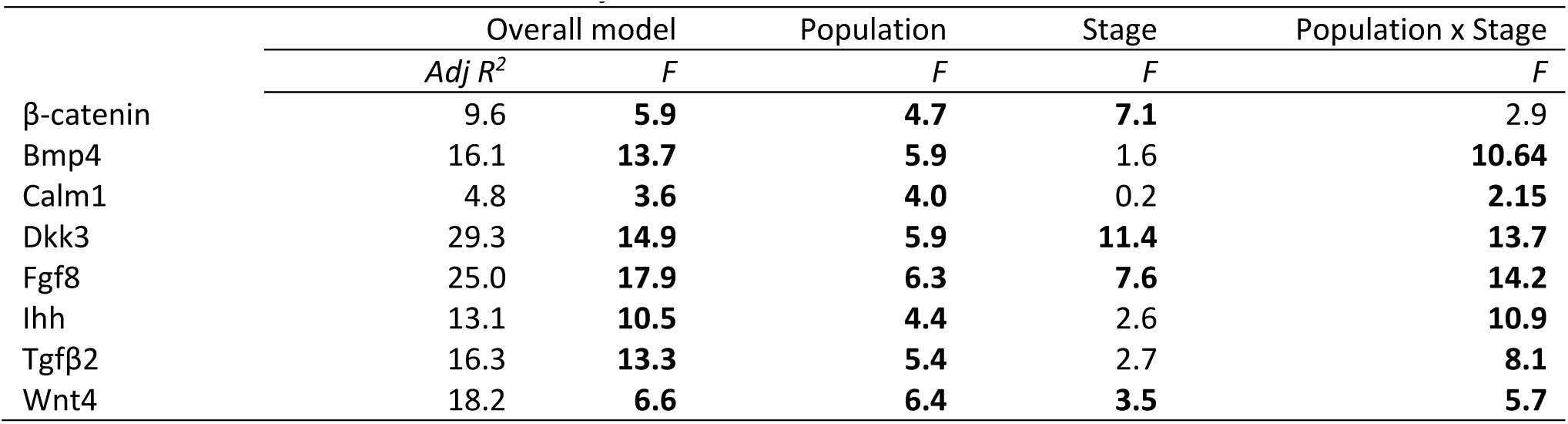
Overall model: effects (Adjusted R^2^) of population and developmental stage on expression of proteins within a group of cells (in each grid box). Bold *F*-values are significant at *P* < 0.05 after within-model Sidak adjustment.

**Supplementary Table S2c.** Overall model for cell shape effects (residuals from Table S2b) on protein expression. Principal components are in Table S2a. Bold *F*-values are significant at *P* < 0.05 after within-model Sidak adjustment.

|  | Cell shape |  | Prin1 | Prin2 | Prin3 |
| --- | --- | --- | --- | --- | --- |
|  | *Adj R<sup>2</sup>* | *F* | *F* | *F* | *F* |
| β-catenin | 19.6 | **15.4** | **26.6** | **6.6** | **17.6** |
| Bmp4 | 11.7 | **17.2** | **26.5** | **5.0** | 1.2 |
| Calm1 | 3.6 | **3.1** | **2.8** | 0.2 | 0.1 |
| Dkk3 | 16.6 | **13.4** | **13.8** | **2.9** | 0.4 |
| Fgf8 | 9.2 | **16.2** | **10.6** | **29.4** | **36.9** |
| Ihh | 5.5 | **8.2** | **20.2** | **6.6** | 0.4 |
| Tgfβ2 | 7.4 | **9.4** | **13.7** | **7.0** | **9.5** |
| Wnt4 | 4.9 | **5.7** | **23.0** | 0.6 | 0.3 |

**Supplementary Table S3a.** Principal component analysis of variation in cell morphology based on *n* = 5,886 cell groups (grid boxes) of *n* = 724,465 mesenchymal cells. Bold values show the largest projection on the original variable. Prin1 mostly captures uniformity in cell circularity variability, Prin2 – uniformity in cell size, and Prin3 – uniformity in cell elongation.

| Variable | Prin1<br>*circularity var* | Prin2<br>*size var* | Prin3<br>*elongation var* |
| --- | --- | --- | --- |
| Area_SD | 0.13 | **0.71** | 0.08 |
| Shape_SD | **0.54** | -0.1 | -0.02 |
| Perimeter_SD | 0.28 | **0.63** | -0.04 |
| Aspect Ratio (AR) _SD | 0.35 | -0.18 | **0.87** |
| Circularity_SD | **0.52** | -0.15 | -0.15 |
| Solidity_SD | 0.46 | -0.20 | **-0.46** |
| Eigenvalue | 3.20 | 1.82 | 0.72 |
| Variance explained | 53.3% | 30.3% | 11.9% |

**Supplementary Table S3b.** Overall model for cell uniformity effects on protein expression (residuals from Table S2b). Principal components are in Table S3a. Bold *F*-values are significant at *P* < 0.05 after within-model Sidak adjustment.

|  | Cell uniformity |  | Prin1 | Prin2 | Prin3 |
| --- | --- | --- | --- | --- | --- |
| | $Adj R^2$ | $F$ | $F$ | $F$ | $F$ |
| β-catenin | 10.1 | **18.0** | **9.2** | **90.7** | **2.0** |
| Bmp4 | 13.7 | **18.3** | **12.9** | **69.1** | 0.1 |
| Calm1 | 3.5 | **5.2** | 0.4 | 2.7 | 0.1 |
| Dkk3 | 16.1 | **9.7** | **5.9** | **12.4** | 2.2 |
| Fgf8 | 7.8 | **13.1** | 1.1 | **82.3** | 0.9 |
| Ihh | 3.0 | **5.0** | **14.3** | **15.0** | 0.8 |
| Tgfb2 | 6.4 | **8.0** | **13.7** | 1.5 | 1.68 |
| Wnt4 | 5.0 | **7.9** | **8.6** | 2.5 | 0.2 |

**Supplementary Table S4.** Least squared means (from mixed effects model) for jammed and unjammed cell states, relative difference between these least squared means ((jammed-unjammed)/unjammed)x100% and analysis of Cell Jamming State on residual protein expression (from Table S2b).

| Protein | Least squared means |  |  | Cell jamming state |  |
| --- | --- | --- | --- | --- | --- |
| | Unjammed | Jammed | Difference, % | $Adj R^2$ | $F$ |
| β-catenin | 17.40 | 25.02 | **+43.81\*** | 10.90 | **9.79** |
| Bmp4 | 4.14 | 0.85 | **-79.50\*\*** | 15.10 | **6.20** |
| CaM | 1.01 | 1.53 | **+51.81\*** | 11.50 | **4.38** |
| Dkk3 | 3.92 | 0.43 | **-89.03\*\*** | 16.97 | **11.27** |
| Fgf8 | 3.60 | 5.53 | **+53.70\*\*** | 13.51 | **6.92** |
| Ihh | 2.48 | 2.00 | -19.30 | 3.10 | 0.93 |
| TGFβ2 | 8.26 | 7.13 | -13.70 | 2.07 | 0.64 |
| Wnt4 | 2.18 | 2.96 | **+35.75\*** | 7.51 | **3.53** |

**Table S5.** Matrices of partial correlations in protein expression in a) unjammed cell state (*n* = 2,212 cell groups), b) jammed cell state (*n* = 690 cell groups), and c) divergence matrix for expression differences between the states. Bold values show correlations significant at *P* < 0.1 in a) and b) and values above average divergence (0.378) in c). Values below c) show contribution of variables to divergence matrix

| a) Unjammed cell state |  |  |  |  |  |  |  |  |
| --- | --- | --- | --- | --- | --- | --- | --- | --- |
|  | Bmp4 | Cam | Bcat | Dkk3 | Fgf8 | Ihh | Tgfb2 | Wnt4 |
| Bmp4 | 1 |  |  |  |  |  |  |  |
| Cam | 0.27 | 1 |  |  |  |  |  |  |
| Bcat | -0.49 | 0.68 | 1 |  |  |  |  |  |
| Dkk3 | 0.68 | 0.43 | 0.44 | 1 |  |  |  |  |
| Fgf8 | -0.63 | 0.29 | 0.57 | 0.45 | 1 |  |  |  |
| Ihh | 0.29 | 0.88 | 0.19 | 0.38 | 0.30 | 1 |  |  |
| Tgfb2 | 0.41 | 0.13 | 0.31 | 0.64 | 0.09 | 0.17 | 1 |  |
| Wnt4 | 0.50 | 0.47 | 0.42 | 0.53 | 0.08 | 0.37 | 0.68 | 1 |
| b) Jammed cell state |  |  |  |  |  |  |  |  |
|  | Bmp4 | Cam | Bcat | Dkk3 | Fgf8 | Ihh | Tgfb2 | Wnt4 |
| Bmp4 | 1 |  |  |  |  |  |  |  |
| Cam | 0.43 | 1 |  |  |  |  |  |  |
| Bcat | -0.09 | 0.08 | 1 |  |  |  |  |  |
| Dkk3 | 0.01 | 0.68 | 0.01 | 1 |  |  |  |  |
| Fgf8 | 0.17 | 0.65 | 0.58 | 0.07 | 1 |  |  |  |
| Ihh | 0.23 | 0.34 | 0.78 | 0.30 | 0.64 | 1 |  |  |
| Tgfb2 | 0.07 | 0.53 | 0.26 | 0.12 | 0.09 | 0.38 | 1 |  |
| Wnt4 | 0.24 | -0.53 | -0.13 | 0.04 | 0.78 | 0.39 | 0.62 | 1 |
| c) Divergence matrix |  |  |  |  |  |  |  |  |
|  | Bmp4 | Cam | Bcat | Dkk3 | Fgf8 | Ihh | Tgfb2 | Wnt4 |
| Bmp4 | 0 |  |  |  |  |  |  |  |
| Cam | 0.16 | 0 |  |  |  |  |  |  |
| Bcat | 0.40 | 0.60 | 0 |  |  |  |  |  |
| Dkk3 | 0.67 | 0.25 | 0.43 | 0 |  |  |  |  |
| Fgf8 | 0.80 | 0.37 | 0.01 | 0.38 | 0 |  |  |  |
| Ihh | 0.06 | 0.54 | 0.59 | 0.08 | 0.34 | 0 |  |  |
| Tgfb2 | 0.34 | 0.40 | 0.05 | 0.52 | 0.00 | 0.21 | 0 |  |
| Wnt4 | 0.26 | 1.00 | 0.55 | 0.49 | 0.70 | 0.02 | 0.06 | 0 |
| contribution: | 1.345 | 1.660 | 1.315 | 1.410 | 1.300 | 0.920 | 0.790 | 1.540 |

**Table S6.**
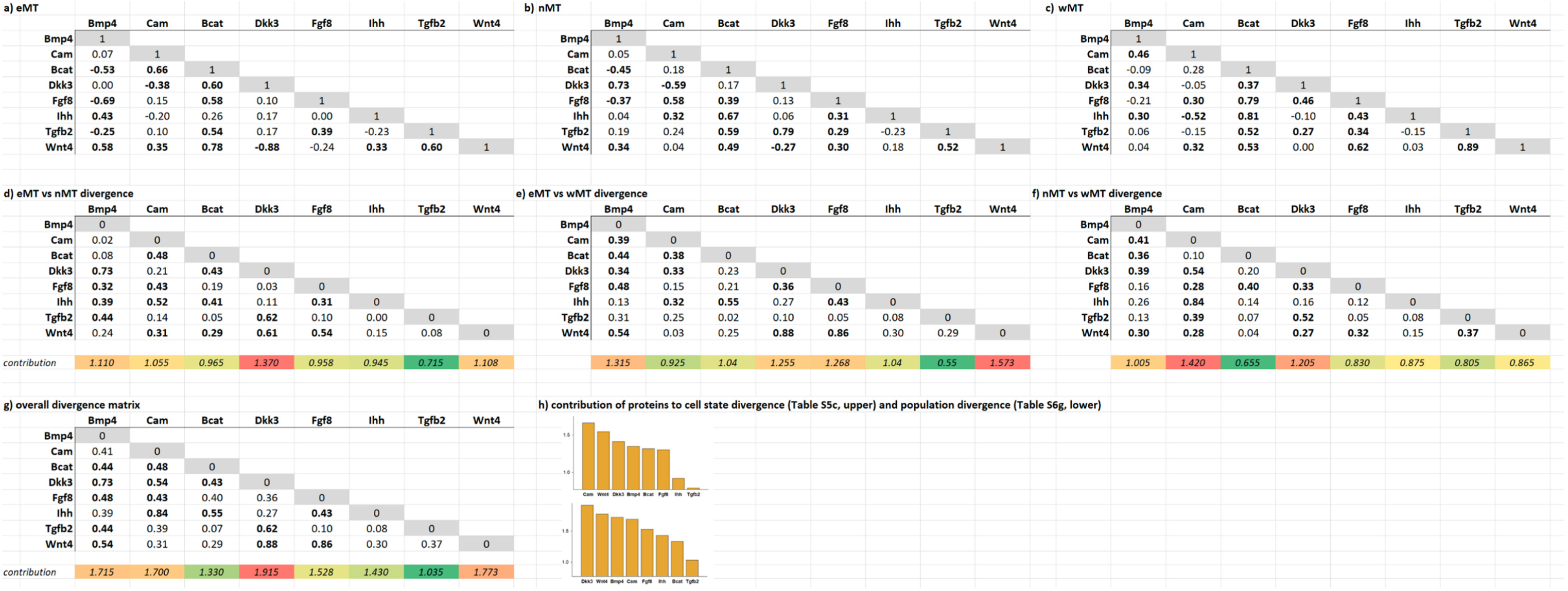
Matrices of partial correlations in protein expression in a) eMT (*n* =131), b) nMT (n=67), and c) wMT (n=107) populations, and pairwise divergence matrices d) eMT vs nMT, e) eMT vs wMT, f)nMT vs w MT and g) overall population divergence matrix. Bold values show correlations significant at *P* < 0.1 in a)-c) and values above matrix-average divergence in d-g (0.25,0.32,0.27,0.43 correspondingly). Values below divergence matrices show contribution of variables to divergence

